# A novel approach for simultaneous detection of structural and single-nucleotide variants based on a combination of chromosome conformation capture and exome sequencing

**DOI:** 10.1101/2024.01.26.577292

**Authors:** Maria Gridina, Timofey Lagunov, Polina Belokopytova, Nikita Torgunakov, Miroslav Nuriddinov, Artem Nurislamov, Lyudmila P Nazarenko, Anna A Kashevarova, Maria E Lopatkina, Elena O Belyaeva, Olga A Salyukova, Aleksandr D Cheremnykh, Natalia N. Suhanova, Marina E Minzhenkova, Zhanna G Markova, Nina A. Demina, Yana Stepanchuk, Anna Khabarova, Alexandra Yan, Emil Valeev, Galina Koksharova, Elena V Grigor’eva, Natalia Kokh, Tatiana Lukjanova, Yulia Maximova, Elizaveta Musatova, Elena Shabanova, Andrey Kechin, Evgeniy Khrapov, Uliana Boyarskih, Oxana Ryzhkova, Maria Suntsova, Alina Matrosova, Mikhail Karoli, Andrey Manakhov, Maxim Filipenko, Evgeny Rogaev, Nadezhda V Shilova, Igor N Lebedev, Veniamin Fishman

**Author notes:** correspondence should be sent to Veniamin Fishman and Maria Gridina.

## Abstract

Effective molecular diagnosis of congenital diseases hinges on comprehensive genomic analysis, traditionally reliant on various methodologies specific to each variant type—whole exome or genome sequencing for single nucleotide variants (SNVs), array CGH for copy-number variants (CNVs), and microscopy for structural variants (SVs). We introduce a novel, integrative approach combining exome sequencing with chromosome conformation capture, termed Exo-C. This method enables the concurrent identification of SNVs in clinically relevant genes and SVs across the genome and allows analysis of heterozygous and mosaic carriers. Enhanced with targeted long-read sequencing, Exo-C evolves into a cost-efficient solution capable of resolving complex SVs at base-pair accuracy. Through several case studies, we demonstrate how Exo-C’s multifaceted application can effectively uncover diverse causative variants and elucidate disease mechanisms in patients with rare disorders.

## Main

The genetic component is an essential factor in most health conditions. In particular, genetic variants dominate the field of rare diseases. The most advanced approaches to human genome analysis, such as whole-genome long-read sequencing, are costly and thus can not be broadly introduced in clinical practice. Routine genetic diagnosis methods include molecular cytogenetic analysis (such as array CGH), karyotyping, and exome sequencing. However, each of these methods is limited in resolution and spectrum of detectable variant types. For example, array CGH (aCGH) is a preferred method for the detection of copy number variants (CNVs), but fails to detect balanced structural variants (SVs) and single-nucleotide variants (SNVs) or short insertions and deletions (INDELs). Whole exome sequencing (WES) is powerful in detection of exonic SNVs, yet can not identify SVs with breakpoints outside exome. Therefore, to choose an appropriate method of genomic diagnosis, it is required to predict the type of variant based on the patient’s phenotype. This challenge asks for a very high qualification, and even for an experienced specialist it is not always possible to guess the genetic etiology of a disease based on the patient’s phenotype. Thus, there is a strong demand for novel methods of genetic assays with reasonable price and sensitivity to detect broad range of genomic variants. To address this challenge, we present a novel method based on chromatin conformation capture that can be used to simultaneously detect structural variants and point mutations in the human genome.

## Results

### Development of the Exo-C: a chromosome conformation capture assay with enriched exome representation

Recently, chromosome conformation capture techniques such as Hi-C were utilized to detect structural variants in the genomes of patients with hereditary diseases^1^. We propose several modifications to this method, allowing to achieve high and uniform coverage of exome sequences while retaining information about chromatin contacts genome-wide (Fig. 1, a,b). The most essential protocol modifications compared to conventional Hi-C method^2^ include using an improved version of DNase I or S1 Hi-C^3,4^ and introducing an exome enrichment step based on a common clinical exome panel (Methods). We designed the resulting protocol as Exo-C, **<u>Exo</u>**me-captured Hi-**C** analysis.

**Figure 1.**
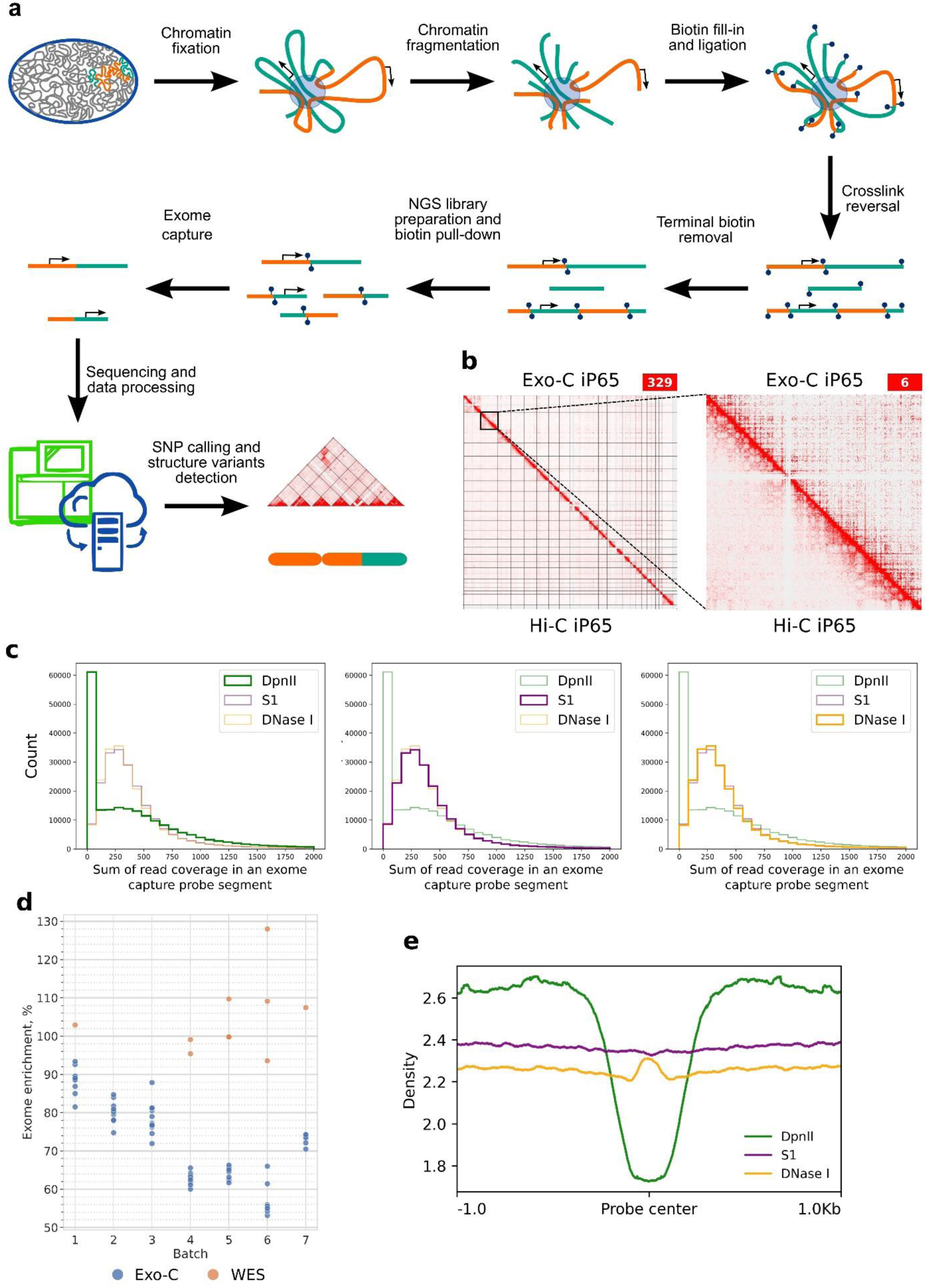
Exo-C achieves high and uniform coverage of exome sequences while retaining information about chromatin contacts genome-wide. **a** – Exo-C Experiment Scheme. The process begins with chromatin fixation and isolation, akin to conventional 3C methods. Subsequently, the chromatin is digested using sequence-agnostic enzyme (DNase I or S1 nuclease). After proximity ligation, products are first selected to contain ligation junctions and then exonic sequences are enriched using hybridisation-based target capture panel. The resulting sequences facilitate the identification of structural variations (SVs) across the genome, single nucleotide variations (SNVs) in the exome, and alterations in spatial contacts of promoters. **b** – Representative interaction heatmap of iP65 cells obtained using Exo-C protocol (above the diagonal line) and S1 Hi-C protocol (below the diagonal line). Red lines correspond to the enriched exonic sequences. **c** – Exome-wide histograms of read coverage depth. Sums of read coverage depth are calculated for segments of all exome capture probes. **d** – Comparative analysis of exome coverage enrichment in Exo-C (blue) and Whole Exome Sequencing (WES, orange) across various batches. Each data point represents an individual sample. The Y-axis quantifies the enrichment ratio, defined as the average coverage within the exome relative to the average coverage outside of the exome. **e** – Read density across segments of exome capture probes, which do not intersect with DpnII restriction sites. The center of the plot is the center of each selected exome capture probe region.

Compared with conventional Hi-C, Exo-C produces similarly high-quality data (Fig. 1, b; Supplementary Table 1). Expectedly, using S1 or DNase I enzymes allows obtaining much more uniform exome coverage compared to DpnII (Fig. 1, c,e). Moreover, Exo-C results in the exome enrichment level only slightly below conventional exome capture data, obtained using the same enrichment panel (Fig. 1, d). These results were reproduced by applying Exo-C to fresh blood samples, cultured fibroblasts or iPS cells (Supplementary Table 1). Finally, comparative analysis of SNVs called from various K562 short-read sequencing datasets reveals that Exo-C performs comparably in SNV detection to other established methods (Supplementary Table 9). Thus, combining chromosome conformation capture with exome enrichment allows obtaining high-quality data to study SNV in exome and chromatin contacts genome-wide.

### Exo-C profiling of 66 human samples and 750 computational models of chromosomal rearrangements

To study the efficiency of the Exo-C we applied it to a cohort of 66 human samples, comprising 28 females, 37 males, and a K562 cell line sample. Prior to Exo-C analysis, the majority of these samples underwent specific molecular and/or cytogenetic profiling (refer to Fig. 2, a and Supplementary Table 1). The cohort encompassed a diverse array of genetic anomalies: 23 balanced translocations, 6 inversions (excluding clinically irrelevant pericentric inversions in heterochromatic regions, undetectable by short-read NGS), 86 CNVs, and several exonic SNVs of clinical significance (detailed in Fig. 2, a and Supplementary Table 1). Additionally, the study included 6 healthy donors with normal karyotype and K562 cells as control subjects. A subset of the probands presented complex SVs, previously characterized by FISH and conventional karyotyping (see below). The aggregated Exo-C dataset that includes all samples, derived from approximately 2 billion reads, incorporates about 1 billion Hi-C contacts, making it one of the most extensive human capture Hi-C datasets currently available.

**Figure 2.**
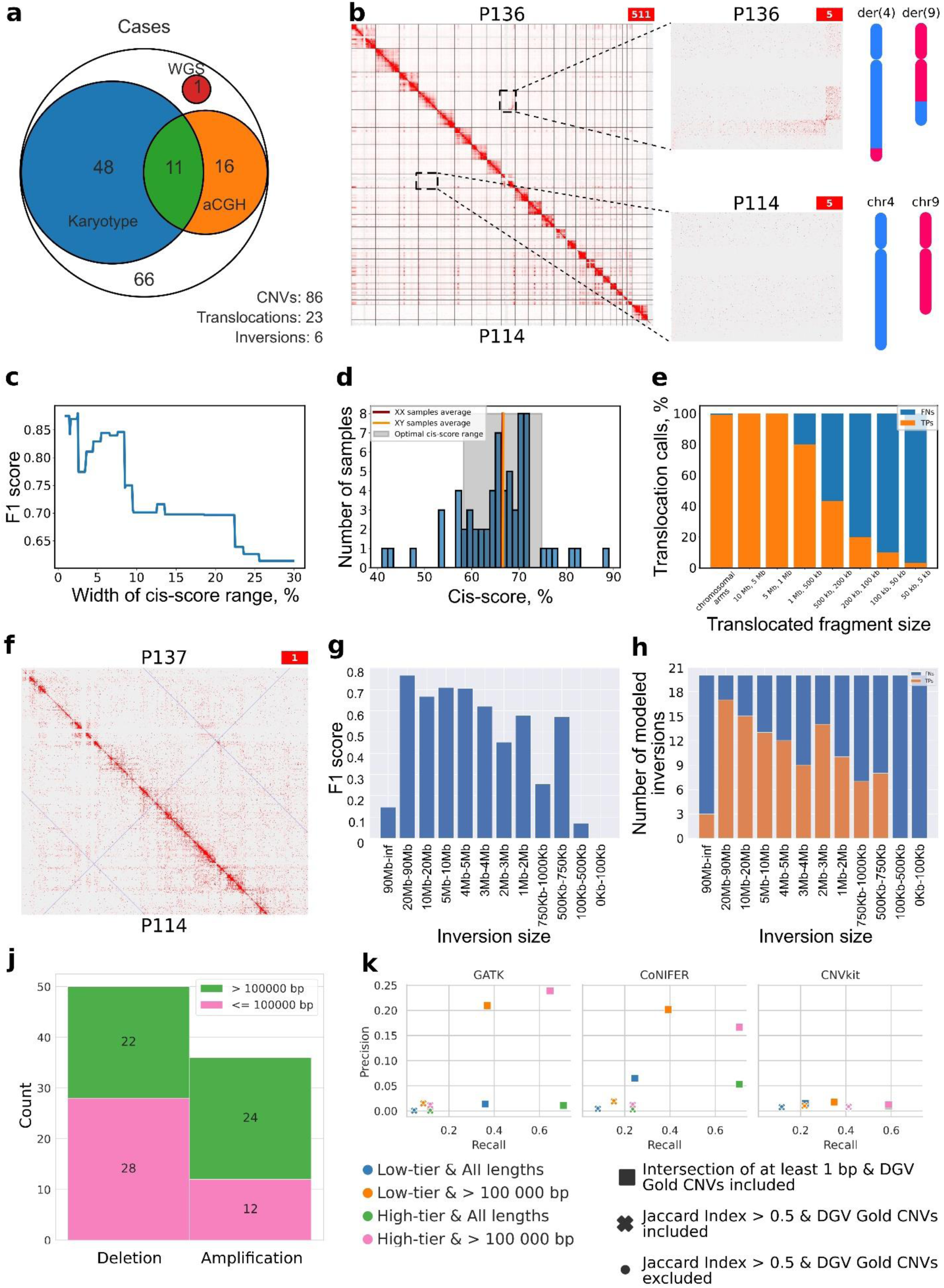
Benchmarking Exo-C against conventional and molecular methods of karyotype and genome analysis (namely karyotyping, aCGH or WGS). **a** – Graphical summary of the 66 samples analyzed in this study. Each sample was examined using the Exo-C protocol, with most undergoing additional orthogonal profiling methods. **b** – Representative example of translocation detected using Exo-C: contacts map of P136 sample P136, which exhibits a chr4-chr9 translocation (above diagonal), and P114, with a normal karyotype (below diagonal). A schematic representation of the translocation and its breakpoint coordinates ([hg19] chr4:174,328,001-174,336,000; chr9:108,424,001-108,432,000) is also included. **c** – Performance of the translocation caller. This graph displays the F1 score, a measure of the translocation caller’s performance. Each data point corresponds to a subset of samples, filtered according to their deviation from the cohort average cis-score. The X-axis specifies the maximum allowable deviation for each data point, while the Y-axis represents the observed F1 score for the corresponding filtered sample set. **d** – The distribution of cis-scores across samples is shown, with vertical lines indicating the average for XX and XY genotypes. The shaded area highlights the optimal cis-score range (±8% from average). **e** – Percentage of True Positive (TP) and False Negatives (FN) calls for modeled translocations of varying lengths. Red bars denote TP calls, while blue bars represent FN. The Y-axis shows the percentage of TP and FN calls for modeled translocations of varying lengths. **f** – Representative example of inversion detected by Exo-C: contacts map of P137 sample (with an inversion on chromosome 7; above diagonal) and P114 sample (with normal karyotype; below diagonal). **g** – F1-score of the inversion-calling algorithm, based on a dataset of simulated inversions of different lengths. **h** – The number of TP calls for simulated inversions, grouped by length intervals, each containing 20 simulations. **j** – Distributions of CNV types and length classes in the low-confidence list of CNVs present in analyzed samples. Bars represent the count of amplifications or deletions with range above (green) or below (pink) 100 000 bp. **k** – Precision and recall of CNV predictions by GATK, CoNIFER, CNVkit tools. Color of markers represent evaluation of tools based on low-or high-confidence lists of CNVs with or without CNV size filtration; shape of markers represents evaluation of tools with or without predictions, intersected with known CNVs for less than 50% (in terms of Jaccard Index) and with or without common CNVs (DGV Gold database). Note that precision and recall outcomes in analyzes excluding or including DGV Gold CNVs are almost similar, therefore corresponding markers may not be fully visible due to overlap.

In this study, balanced SVs were initially identified via microscopy-based methods, typically involving rearranged fragments exceeding several megabases in size and the breakpoints were defined at a resolution of cytobands (detailed in Supplementary Table 1). To evaluate the Exo-C protocol’s effectiveness at finer resolutions, we expanded our analysis to include a range of SV sizes. This was achieved by simulating Exo-C maps for SVs of varying dimensions, utilizing the Charm^5^ framework for *in silico* generation.

Examination of Exo-C maps readily revealed megabase-scale SVs, as evidenced in figure 2, b. Despite a pronounced bias towards exome sequences, these large-scale SVs induce marked alterations in chromatin contact patterns across entire chromosomes. In contrast to WGS data, where only reads adjacent to the breakpoint are indicative of SVs, Exo-C maps demonstrate altered contact patterns at loci situated millions of base pairs from the breakpoint, thereby signifying the presence of SVs (refer to Fig. 2, b).

Despite being visually distinguishable, SVs were not called using the state-of-the-art chromosome conformation capture data analysis tool such as EagleC (F1-score for translocations equals to 0.15, F1-score for inversions equals to 0.03, F1-score for CNV equals to 0 since no CNV were reported). To perform unbiased, automatic calling of chromosomal rearrangements and allow detection of smaller SVs that are not always visible on whole-genome Exo-C maps, we developed dedicated computational algorithms detecting translocations and inversions, and benchmarked existing CNV-detecting tools (Methods).

### Automatic detection of chromosomal translocations

Our methodology demonstrates robust precision and recall in human samples for translocation calling, as shown in Figure 2, c. In patient samples previously analyzed by conventional karyotyping, our method achieved a Precision of 36%, Recall of 100%, with 25 True Positive (TP) calls, 45 False Positive (FP) calls, and no False Negatives (FN). We observed that the quality of experimental data, quantified by the cis-to-trans contacts ratio, significantly influences the caller’s performance. Higher precision is noted in samples with metrics akin to control samples (Fig. 2, c,d).

Exo-C map for each FP call was examined visually, and for 12 FP calls we observed a clear translocation pattern (Supplement Figure 1; Supplement Table 2). Given that some predicted insertion sizes fell below the detection limit of microscopy, (the smallest detected by our algorithm was ∼20 kb, Supplementary Figure 1), it is plausible that these FP calls might represent genuine SVs, undetected by conventional diagnostic techniques. Importantly, in many cases translocation breakpoints were resolved at high resolution, allowing primer design for PCR-validation of the breakpoints (see examples below). For breakpoints where we were not able to design primers, we performed FISH with insertion-specific probes. This approach validated 5 out of 6 tested FP translocation calls using orthogonal methods (PCR or Spectral Karyotyping), as elaborated in the ‘Cases’ section.

Upon refining our analysis by filtering 17 samples based on their ‘cis/trans’ ratio (58% < Cis < 75%; Fig. 2, c,d) and reclassifying the validated translocations as True Positives, the revised metrics showed a Precision of 73%, Recall of 100%, with 22 TP, 8 False Positives (FP), and no False Negatives (FN). It’s important to note that this efficiency may be an underestimation, as some FP calls could not be further verified due to the unavailability of additional sample material for orthogonal testing.

To investigate the impact of translocated segment size on method sensitivity, we employed the Charm *in silico* modeling framework to create Exo-C contact maps for 330 simulated translocations or insertions, with sizes ranging from 10 kb to 140 Mb (detailed in Methods). As anticipated, the recall rate of translocations increased with the size of the insertions (Fig. 2, e). Detection rates increased from approximately 5% for the smallest insertions (5-50 kb) to around 45% for larger events (200-500 kb), achieving 100% sensitivity for fragments exceeding 1 Mb. The overall F1-score for all modeled cases remained 0.77, with a Precision of 87%, Recall of 68%, comprising 226 true positives, 34 false positives, and 104 false negatives.

### Automatic detection of inversions

Inversions, like translocations, manifest as distinct contact pattern changes on Hi-C maps (Fig. 2, f). Of the six inversion cases analyzed, our method successfully detected three, alongside 220 false positive inversions, yielding a precision of 10% and a recall of 50%. Notably, the sensitivity of inversion detection is closely tied to the quality of experimental data, quantified by the cis-to-trans contacts ratio, similar to translocation calling. Remarkably, the false positive count significantly reduces from 220 to 33 when applying the same cis-to-trans filter to the samples, while the recall remains consistent at 50% for this subset. Further investigation reveals that some false-positive calls correspond to inversions undetectable by cytogenetic analysis but verifiable through alternative methods. For instance, a 2 Mb inversion identified in P10 data was initially reported as a false positive. Subsequent validation using long-read sequencing, as detailed below, confirmed this inversion and its correlation with the observed phenotype in the P10 case. Consequently, the actual precision of the inversion detection algorithm is likely higher than initially estimated, suggesting an overestimation of false positive calls.

We next assessed the algorithm’s performance across a range of inversion sizes, from 100 kb to several megabases, utilizing 240 *in silico* generated inversions (Fig. 2, g,h). This benchmarking, using the series of simulated Exo-C maps, yielded an average F1-score of 0.49 for all inversion lengths (Precision: 56%, Recall: 45%, True Positives: 107, False Positives: 85, False Negatives: 133). The analysis revealed a positive correlation between the algorithm’s performance and the size of the inversion, with larger inversions yielding higher scores. However, it was noted that the algorithm encounters challenges in detecting inversions exceeding 90 Mb, primarily due to their breakpoints being situated at the chromosomal ends, which complicates the recognition of inversion patterns on Hi-C maps.

### Detection of CNVs

Copy-number variants (CNVs) modify the representation of genomic loci in NGS datasets, enabling their identification through analysis of read coverage distribution. We hypothesized that Exo-C results should allow analysis of genomic coverage similar to the whole-exome data. To evaluate this, we tested existing CNV prediction tools on Exo-C datasets. Utilizing available aCGH data, we compiled a list of CNVs, which we further categorized into high-confidence (clinically relevant CNVs validated by qPCR or other methods) and low-confidence groups (identified in aCGH data but lacking orthogonal confirmation) (detailed in Methods; Supplementary Figure 2, a and Supplementary Table 3). This categorization resulted in 46 CNVs in the high-confidence list and 86 in the low-confidence list. The distribution of these CNV types and their length classes is illustrated in figure 2, j.

Leveraging the compiled dataset alongside corresponding Exo-C data, we conducted a benchmarking study of three CNV callers — CNVkit, CoNIFER, and GATK — originally designed for WES data analysis. Despite not being tailored for Exo-C data, these tools demonstrated the capacity to identify over 60% of CNVs in our samples (square markers in figure 2, k; Supplementary Table 4). However, the overlap between CNVs detected by these tools and those reported by aCGH was often partial. When predictions were filtered to include only those with a Jaccard Index greater than 50% (cross markers in figure 2, k), the performance of GATK and CoNIFER, in both precision and recall, significantly decreased. This filtration method only marginally impacted CNVkit’s performance, possibly due to its inherently lower precision prior to any filtration.

Implementing an additional filtration layer, which excludes CNVs commonly found in the population as per the DGV Gold database, significantly reduces the number of false positive predictions. This effect is particularly evident in the GATK results, where the count of small CNV predictions (less than 100,000 bp) is reduced by nearly 70%. A similar, albeit less marked, reduction is observed across all evaluated tools, both for various CNV tiers and size-based filtration (as detailed in Supplementary Table 4). However, the overall precision of these tools, initially modest, does not exhibit a substantial absolute increase post-filtration (indicated by dot markers in figure 2, k, closely aligning with cross markers).

We hypothesized that CNVs from the high-confidence tier of clinically-relevant variants would exhibit more comprehensive detection, given their lower likelihood of being common population variants. Corresponding results affirmed this hypothesis, revealing that high-confidence tier CNVs were indeed predicted with greater recall across all assessed tools (Fig. 2, k). The precision for high-confidence CNVs surpassed that of the low-confidence tier exclusively in the case of GATK.

### Exo-C can be employed to detect mosaic translocation carriers

Exo-C demonstrates exceptional precision and recall in detecting translocations, particularly excelling in cases involving large, megabase-scale translocations. This suggests a potential for detecting SVs even in a minor subset of cells, a capability crucial for applications in cancer research or germ-line mosaicism studies. To investigate this potential, we employed our translocation calling methods in both *in silico* and *in vitro* experiments simulating ‘mosaic cell populations’. In the *in silico* experiment, we merged random reads from pairs of samples, one carrying known structural variants and the other with a normal karyotype, in varying proportions (as detailed in Supplementary Table 5).

We proportionally adjusted the data inputs so that 10% to 90% of the reads in the merged dataset originated from the sample with translocations. As depicted in Figure 3A, our method can reliably identify heterozygous translocations in samples with a fraction above 80%, and some heterozygous translocations are detectable when the mix contains only 40% of data from the translocation-carrying sample. It is important to note that, since we mixed data from heterozygous translocation carriers, the actual fraction of alleles with the structural variant is half of the indicated data fraction.

**Figure 3.**
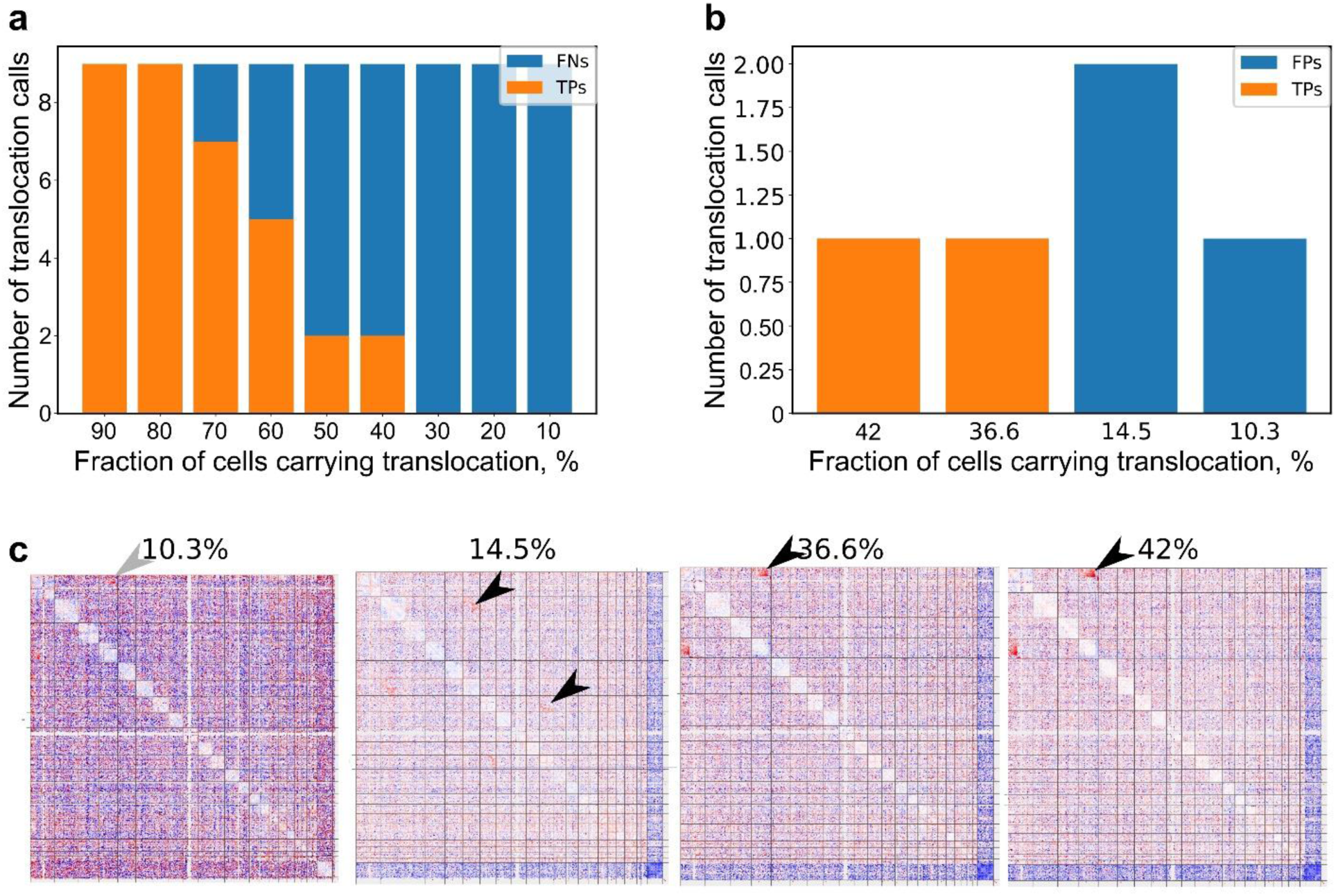
Detecting mosaic translocation carriers using Exo-C. a and **b** – Recall of the translocation caller. Y-axis show number of TP calls for samples with different fraction of translocation-carrying cells (%) for *in silico* (**a**) and *in vitro* experiments (**b**). Red bars indicate True Positive calls, blue bars – False Negative. **c** – Representative interaction Exo-C heatmap (Log(Observed/Control); Observed – mixed samples; Control – patient without translocations). Black arrows indicate visually detectable chromosomal rearrangements, gray arrows – expected rearrangements.

In the *in vitro* experiment, we generated cell mixtures from P62+P114 samples (where P62 exhibits t(1;4) and P114 has a normal karyotype) with 10.3%, 36.6%, and 42% of cells carrying the translocation; and from P69+P114 samples (P69 with t(2;6) and t(7;11), P114 normal) with 14.5% translocation-carrying cells. The exact proportion of cells with chromosomal rearrangements was determined based on the SNV analysis. Applying Exo-C to these samples corroborated previously described findings (Fig. 3, b). Although our bioinformatic tool did not detect translocations in samples with less than 36% SV-carrying cells, visual analysis suggests that the translocation pattern is still discernible at a 14.5% fraction (Fig. 3, c). These observations lead us to speculate that further refinement of computational tools could enhance the precision of translocation detection in mosaic cell populations.

### Augmenting Exo-C with targeted Oxford Nanopore sequencing allows cost-efficient detection of structural variants with base-pair resolution

Although Exo-C shows high sensitivity in SVs detection, variants can not be resolved with single-nucleotide resolution, and in some cases resolution is too low to design primers for breakpoints amplification and sequencing. Yet, precision of the Exo-C method is high enough for using cost-efficient targeted sequencing of the detected translocation breakpoints. To benchmark this approach, we selected 5 cases involving 23 Exo-C-identified breakpoints, with 17 confirmed via orthogonal techniques. Only two junctions were validated by PCR with single-nucleotide resolution, others were confirmed by karyotyping (Supplementary Table 6). We applied whole-genome nanopore sequencing (WGS), adaptive sampling (AS) and modified (see Methods) nanopore Cas9-targeted sequencing (nCATS)^6^ approaches to this cohort (Fig. 4, a,b). We confirm that nCATS greatly improves coverage of the breakpoint regions (Fig. 4, c), whereas substantially deeper sequencing would be required to achieve the same breakpoint coverage without enrichment. The adaptive sampling approach, while displaying a balanced performance compared to previous methods, offers the advantage of validating breakpoints without requiring specialized library preparation such as the production of sgRNAs for nCATS. It is important to note that although the coverage of target regions in AS is lower compared to nCATS, it presents a viable option for breakpoint validation. For both enrichment and WGS we limit sequencing depth to the same value (up to 1.5X coverage, Supplementary Table 6) to ensure reasonable cost of analysis in future clinical use.

**Figure 4.**
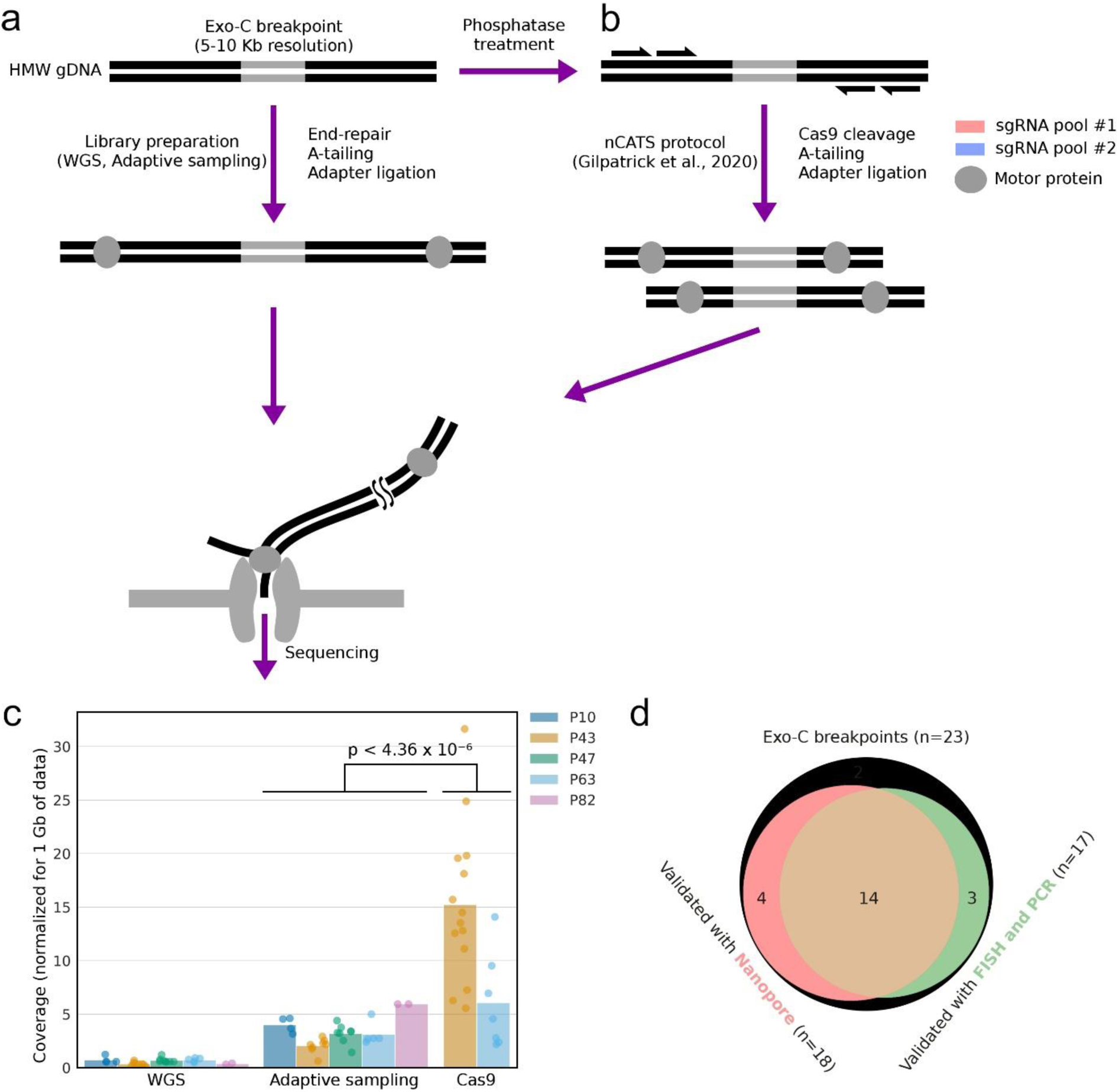
**Combination of Exo-C and nanopore sequencing enables detection of breakpoints with single-nucleotide resolution**. **a** – Assembly of Nanopore ligation libraries for WGS and adaptive sampling. **b** – Enrichment of breakpoint regions with Cas9 utilizing a combination of tiling and nCATS approaches. **c** – Comparison of coverage in breakpoint regions between different Nanopore library types (normalized for 1 Gb of data, coverage calculated for 5 kb bins). **d** – Graphical representation of breakpoints detection by Exo-C, nanopore sequencing, and other methods.

We analyzed sequencing results both manually (aiming to detect any chimeric reads in a vicinity of the Exo-C-derived breakpoints) or automatically using the NanoSV^7^ software. The results demonstrate our ability to validate the majority of translocations identified using Exo-C through nanopore sequencing, achieving single-nucleotide resolution (Fig. 4, d and Supplementary Table 6). In particular, we were able to confirm and resolve at single-nucleotide level 18 out of 23 Exo-C-detected breakpoints (78%) and 4 breakpoints which were not validated previously by orthogonal methods were validated in this assay (Fig. 4, d). As expected, targeted sequencing methods (AS and nCATS) outperformed nanopore WGS in effectively covering breakpoint regions. Automatic breakpoint detection with NanoSV was effectively possible only in regions with high coverage, highlighting the possibility for use in combination with Cas9 enrichment.

Importantly, the targeted sequencing approaches require only 20-30% of the resources of a MinION or GridION flow cell, which allows cost-effective breakpoint verification for less than $300 per sample. This affordability underscores the possibility of implementing targeted nanopore sequencing as a valuable complement to Exo-C, presenting an economical solution for breakpoint validation in clinical applications. These findings emphasize the potential of combining Exo-C with targeted nanopore sequencing as an efficient and budget-friendly tool for clinical use.

### Application of Exo-C Technology in Resolving Clinical Cases

This study encompassed several patients with congenital diseases who had previously eluded molecular diagnosis. We demonstrate how Exo-C can identify rare genomic variants in these cases, and subsequent analyses elucidate their relevance to disease pathogenesis

### Patient P10. Detecting inversion which causes morbid gene disruption

An 8-month-old girl was referred for evaluation because of developmental delay, hypotonia, feeding problems, delayed motor milestones, and abnormal phenotype including microcephaly, almond-shaped palpebral fissures, low-set ears, posteriorly angulated ears, wide nasal bridge, fifth-finger clinodactyly, pes valgus.

Karyotyping revealed a *de novo*, apparently balanced translocation between chromosomes 5 and 10 in the patient: 46,XX,t(5;10)(q11.2;q11.2)dn. aCGH did not detect unbalanced chromosomal rearrangements. Utilizing Exo-C, we resolved the genomic breakpoints of this translocation with high resolution (Fig. 5, a). This resolution enabled the amplification and sequencing of the junction region (Fig. 5, b). The sequencing data revealed that the breakpoint on chromosome 10 intersects the *PCDH15* gene, while the breakpoint on chromosome 5 is situated in an intergenic area proximal to the *RNF180* gene promoter. Intriguingly, neither of these genes appear to correlate with the patient’s clinical phenotype.

**Figure 5.**
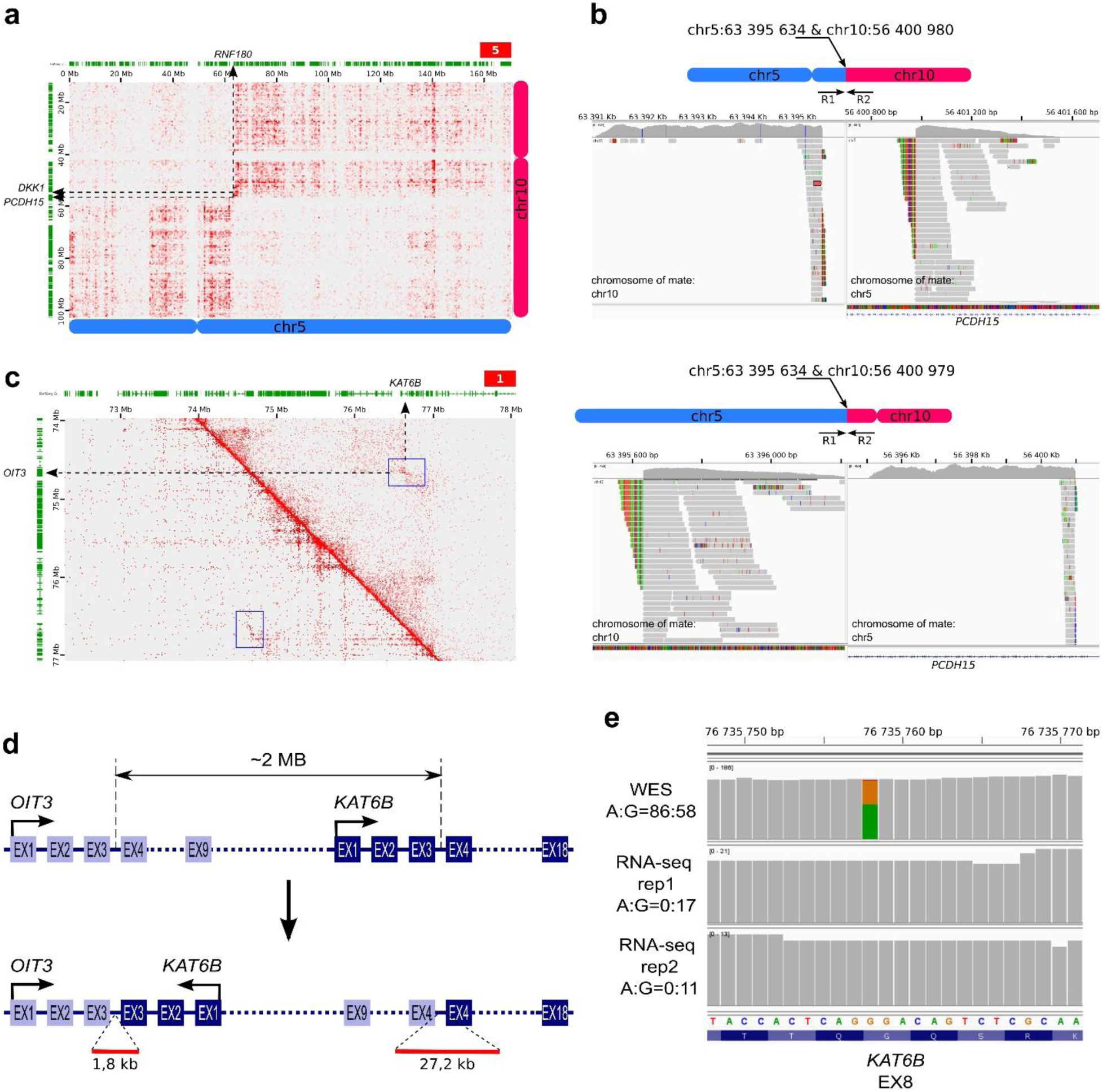
**Detection and functional characterization of balanced translocation and inversion using Exo-C technique**. **a** – An Exo-C contact map indicating balanced translocation pattern for P10 case. **b** – IGV screenshot showing alignment of reads obtained by amplification and sequencing of the t(5;10) junction. **c** – Hi-C (above diagonal) and Exo-C (below diagonal) maps indicating the presence of balanced translocation in P10 genome. **d** – Scheme of the inversion detected using Exo-C and Hi-C data. Genomic breakpoints of inversion between *OIT3* (breakpoint coordinate – chr10:74 666 324) and *KAT6B* (breakpoint coordinate – chr10:76 646 623) were identified by 1 778 bp and 27 189 bp nanopore sequencing reads (red lines), spanning of third introns *OIT3* and *KAT6B*. **e** – IGV screenshot showing allelic representation of single-nucleotide variant in the *KAT6B* exon 8 according to the Exo-C sequencing data and transcriptome sequencing data.

In our analysis of genes proximal to the translocated fragments junction, we identified the *DKK1* gene approximately 1.5 Mb from the chromosome 10 breakpoint (Fig. 5, a). We hypothesized that *DKK1* misregulation might contribute to the disease phenotype. Supporting this, both low-input Hi-C and the original Exo-C data revealed altered chromatin interactions near the translocation breakpoints, affecting the *DKK1* promoter (Fig. 5, a). To test the hypothesis of *DKK1* gene misregulatoin, we derived induced Pluripotent Stem Cells (iPSCs) from the patient’s peripheral blood and analyzed gene expression in these cells and their differentiated primitive streak derivatives (Supplementary Figure 3, a,b). Given the known high expression of *DKK1* in primitive streak cells, we conducted digital PCR analysis on patient-derived iPSС and primitive streak cells. Contrary to our hypothesis, the results indicated no significant difference in *DKK1* expression between the patient and control cells, challenging the hypothesis of *DKK1* misregulation (Supplementary Figure 3, c).

Since translocation between chromosomes 5 and 10 can not explain P10 phenotype, we apply Exo-C variant callers to identify other clinically relevant genomic variants. No pathogenic SNVs were identified; however, the Exo-C inversion caller detected a ∼2 Mb inversion on chromosome 10, situated approximately 20 Mb from the previously identified translocation breakpoints. Given the significant genomic distance between breakpoints, we propose that translocation and inversion occur as two independent mutations on chromosome 10. The inversion, undetected in karyotyping due to its size, was corroborated by low-input Hi-C analysis, which confirmed the inversion pattern (Fig. 5, c). Additionally, the presence of the inversion was supported by split-reads identified in nanopore sequencing data (see above). Based on these data, inversion breakpoints can be mapped within the 3rd intron of the *KAT6B* gene and the 3rd intron of the *OIT3* gene, notably disrupting the KAT6B gene (Fig. 5, d).

Loss-of-function variants in the *KAT6B* gene are known to cause SBBYSS syndrome (OMIM #603736) in an autosomal-dominant manner, potentially explaining the patient’s phenotype. To evaluate the hypothesis of *KAT6B* gene disruption, we conducted transcriptome analysis in iPSC. This analysis revealed abnormal splicing products consistent with the predicted inversion breakpoint within the 3rd intron of *KAT6B* (Supplementary Figure 3, d). Furthermore, based on Exo-C data, we identified a benign heterozygous SNV in exon 8 of the *KAT6B* gene (Fig. 5, e). RNA-seq data, however, indicated the expression of only one allele of this gene (Fig. 5, e). These findings collectively confirm that only one functional copy of the *KAT6B* gene is expressed, correlating with the patient’s observed clinical manifestations.

Thus, Exo-C and Hi-C analysis allowed us to characterize two balanced structural variants and develop hypotheses about causative genes implicated in the disease. Pursuing this line of investigation, we were able to characterize inversion affecting the *KAT6B* gene.

### Patient P47

An 8-year-old boy was examined by a geneticist regarding developmental delay (low height and weight) and multiple exostoses. The first exostosis on the rib was noted at birth, the second appeared on the arm at the age 2.5 years. At the time of examination, the patient had multiple exostoses. A dysmorphic features like dolichocephaly, epicanthus and right dysplastic ear were noted.

Conventional karyotyping of G-banded chromosomes revealed a balanced translocation between chromosomes 8, 11, and 21: 46,XY,t(8;11;21)(q23;q22;q21)[10], aCGH did not reveal any unbalanced chromosomal rearrangements.

In Exo-C assay of this case, we did not detect any pathogenic SNVs. However, the Exo-C data analysis confirmed a complex apparently balanced chromosomal rearrangement involving chromosomes 8, 11, and 21. This analysis accurately resolved the breakpoints of these rearrangements and additionally identified two translocations: one between chromosomes 20 and 21, and another involving fragments of chromosomes 11, 8, and 20 (Fig. 6, a,b). These findings were subsequently confirmed using Spectral Karyotyping (SKY) (Fig. 6, c, e) and Oxford Nanopore long-read sequencing. Notably, one breakpoint was located within the *EXT1* gene (Fig. 6, b,e), a known causative gene for autosomal-dominant hereditary multiple exostoses (OMIM #133700). The disruption of this gene elucidates the patient’s clinical phenotype. This case exemplifies how the delineation of translocation breakpoints via Exo-C allows the classification of balanced chromosomal translocation as pathogenic.

**Figure 6.**
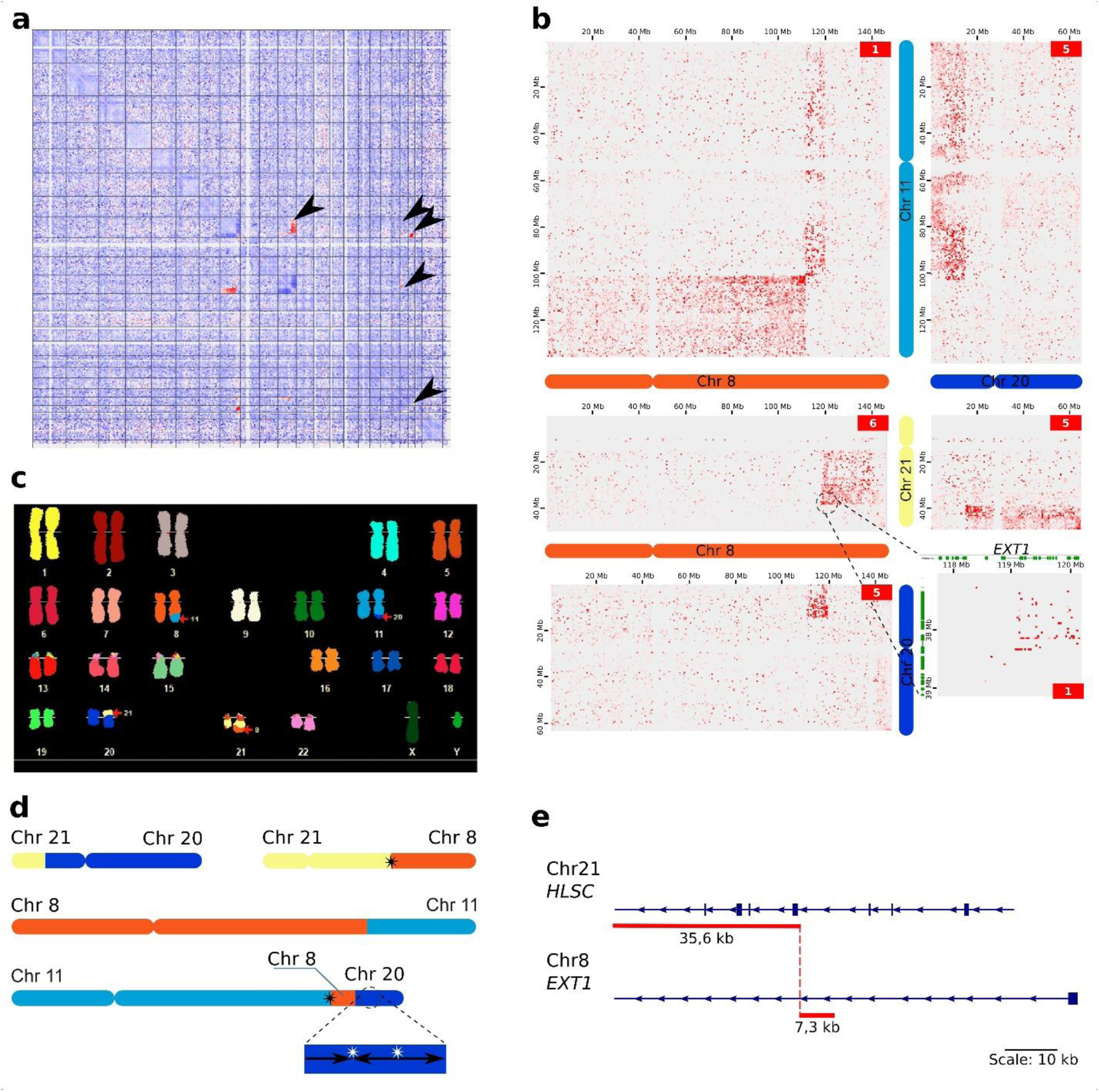
Resolving breakpoints of complex chromosomal rearrangement through Exo-C analysis provides evidence of its pathogenicity. **a** – Exo-C contacts normalized as Log(Observed/Control), where Observed is contact counts for P47 which exhibits complex SV, and control is Contact counts for control sample without translocations. Black arrows indicate Exo-C contacts at translocation breakpoints. **b** – Interchromosomal Exo-C contact maps for P47 sample supporting SV structure. **c** – SKY results supporting SV structure. **d** – Schematic representation of SV detected in P47 cells based on Exo-C analysis. Asterisks indicate breakpoints identified with single-nucleotide resolution breakpoints by ONT sequencing. **e** – Genomic breakpoint of translocation between *EXT1* (breakpoint coordinate – chr8:119 069 497) and *HLSC* (breakpoint coordinate – chr21:38 321 275) was identified by a 41 312 bp sequencing read (red line), spanning 7.3 kb of first intron *EXT1* and 35.6 kb of second intron *HLSC*.

### Patient P123

A 41 years old male was under examination after he has а stroke of the left MCA (middle cerebral artery) of ischemic type, with right-sided hemiparesis and gradual regression of focal disorders. Repeated episode of cerebrovascular accident occurred after six months, with hemiplegia of the left arm and leg, facial asymmetry. CT revealed a hematoma in the right hemisphere measuring 20 cm^3^, against the background of lesions arteriopathy with subcortical infarcts. The consequences of multiple repeated cerebral infarctions of the deep parts of both hemispheres of the white matter of the brain of varying duration were revealed. Neurometabolic therapy was applied with positive effects. Patient previously reported migraine-like headaches since age 20. The patient has a 17-year-old daughter who complains of migraine-like headaches.

While Exo-C analysis did not reveal any pathogenic SVs or CNVs, it identified a SNV rs1555729452 (Cys222Tyr) within the *NOTCH3* gene, a known causative gene for cerebral autosomal dominant arteriopathy with subcortical infarcts and leukoencephalopathy (CADASIL; MIM 125310). Classified as pathogenic based on the American College of Medical Genetics and Genomics (ACMG) guidelines, this variant accounts for the patient’s clinical presentation. This case underscores the capability of Exo-C data to pinpoint pathogenic SNVs within the exome, showcasing a performance comparable to traditional exome sequencing methods.

## Discussion

In this study, we demonstrate the utility of Exo-C, a combination of modified chromosome conformation capture assay and exome enrichment, in elucidating a broad spectrum of clinically significant genomic variants in patients with monogenic and chromosomal diseases. With cost comparable to standard exome sequencing, Exo-C notably enhances it in the detection of balanced translocations and inversions. Thus, Exo-C facilitates the simultaneous identification of SVs and SNVs, surpassing the capabilities of traditional WES.

Complementing the experimental application of Exo-C, this publication introduces a suite of computational tools specifically tailored for Exo-C data analysis, focusing on SV calling. The development of automate callers is crucial for the clinical implementation of this technique as well as for an unbiased evaluation of its performance. Existing computational solutions, primarily designed for genome-wide cancer Hi-C datasets, fall short when applied to Exo-C data. Addressing this gap, we have developed and rigorously benchmarked computational tools that facilitate automated Exo-C-based SV detection, thereby enabling users to process their data with greater efficiency and precision.

While Exo-C significantly enhances standard exome analysis, it is important to acknowledge its limitations. A primary constraint is the diminishing recall of Exo-C SV callers for smaller-sized SVs. The tool exhibits relatively modest performance in identifying translocations and inversions under 100 kb. Intergenic events smaller than 100 kb remain challenging to reliably identify without resorting to impractically high sequencing depths. However, it’s noteworthy that smaller translocations, and potentially inversions, overlapping with exonic sequences are more detectable due to the significantly higher coverage of these fragments compared to the genomic average.

A second limitation of Exo-C concerns CNV detection. The precision and recall of CNV callers on Exo-C data are relatively low, mirroring the performance observed with WES data^8,9^. This is partly because, for CNV calling, Exo-C data analysis was limited to genomic coverage assessment. The current effectiveness could be enhanced by incorporating analyses of specific contact frequency alterations attributable to CNVs. However, insights from studying contact frequency changes induced by inversions suggest that significant improvements in detecting CNVs smaller than 100 kb are unlikely. Consequently, aCGH and low-coverage WGS currently offer a more favorable balance between cost and the quality of CNV detection.

Considering the strengths and limitations of the Exo-C technique, its application can be particularly advantageous in several diagnostic scenarios. Firstly, Exo-C is well-suited for patients with karyotypic abnormalities of undetermined significance detected through microscopy-based methods. Here, Exo-C can precisely resolve breakpoints to an extent that facilitates the identification of genes disrupted by SVs. Additionally, Exo-C is capable of detecting exonic SNVs, testing alternative hypothesis about the nature of genomic variant underlining the patient’s phenotype.

Secondly, Exo-C can serve as an alternative to standard WES for primary screening in patients suspected of harboring disease-causing SNVs. Exo-C can be adapted to various hybridization-based capture panels, enabling custom assays tailored to different patient groups. Exo-C not only delivers the same insights as WES but also identifies submicroscopic SVs within or adjacent to genes of interest.

Thirdly, Exo-C is valuable following aCGH analysis, particularly when a CNV affects one copy of a clinically relevant haplosufficient gene. In such cases, Exo-C can aid in identifying loss-of-function mutations in the second allele of the implicated gene.

Recent advancements in research indicate the potential of chromatin interactions for SNV phasing. This presents another promising application for the Exo-C method: distinguishing between cis and trans configurations in compound heterozygous scenarios. However, realizing this application will necessitate the development of specialized bioinformatic tools tailored for Exo-C data analysis.

Finally, Exo-C offers valuable opportunities to explore the regulatory effects of previously identified SVs, utilizing data on chromatin contact frequencies to gain insights into their functional implications. A unique aspect of Exo-C is its ability to analyze spatial interactions of gene promoters, which are crucial for understanding diseases caused by missregulation of gene expression^10^. While computational predictions of alterations in chromatin contacts exist^11^, their accuracy remains insufficient to replace empirical data^12^. Consequently, the chromatin interaction patterns unveiled by Exo-C offer valuable insights for variant interpretation, bridging a critical gap in genomic analysis

In the latter part of this manuscript, we illustrate this through three use-cases, demonstrating Exo-C’s capability in identifying novel pathogenic variants and guiding the investigation into their pathogenicity.

The integration of Exo-C with other techniques of molecular analysis can significantly enhance its capacities. Here we highlight integration of Exo-C with targeted Oxford Nanopore sequencing, enabling the reconstruction of complex SVs with single base-pair resolution in a cost-effective manner.

In summary, the Exo-C technique, as detailed here, broadens the scope for identifying causative variants in patients with genetic diseases, offering significant potential in genetic diagnostics and research.

## Methods

### Human samples

All samples were collected from authorized medical centers. The study was approved by the local ethics committee of the Institute of Cytology and Genetics (protocol number 17, 16.12.2022) and the local ethics committee of the Tomsk Medical Research Center (protocol number 15, 28.02.2023). An Informed Consent was obtained from all patients or their parents/representatives included in the study. Blood samples were collected as described in^3^. Human cell lines used in this study are described in Supplementary Table 1.

### Exo-C experiment

Exo-C protocol includes a 3C part followed by exome enrichment. The 3C part was performed either following DNase I protocol^3^, or S1 protocol^4^. Briefly, 2,5 million cells were fixed in 1% formaldehyde, lysed, and digested overnight with DNase I or S1 nuclease. Digested DNA ends were marked with biotin-14-dCTP and ligated overnight using T4 DNA ligase. Formaldehyde crosslinking was reversed by incubation with Proteinase K at 65°C overnight, followed by ethanol precipitation. Biotin from unligated ends was removed by T4 polymerase treatment. KAPA HyperPlus kit was used for NGS library preparation. Biotin-filled DNA fragments were pulled down using Dynabeads MyOne Streptavidin C1 beads before library amplification. Exome enrichment was performed using KAPA HyperExome Probes and KAPA HyperCap kit according to the manufacturer’s instructions.

### iPS cells generation and differentiation

Reprogramming of human blood cells into iPS cells was performed by episomal vectors cocktail contained: reprogramming vectors expressed OCT4 (addgene #41813), MYC and LIN28 (addgene #41855), shRNA against p53 (addgene #41856), SOX2 and KLF4 (addgene #41814), EBNA1 (addgene # 41857), according to the previously described method^13^. iPS cells were cultured as described before^14^. Two iPS cell lines from healthy donor^14^ and from P10 were differentiated into primitive streak cells^15^.

### Spectral karyotyping (SKY)

Muticolor FISH analysis for verification of predicted translocations was performed on patient’s metaphase chromosomes using SkyPaint™ DNA Kit H-10 for Human Chromosomes (Applied Spectral Imaging, Israel). Visualisation of hybridization signals was done using Olympus BX53 microscope (Olympus Life-science) equipped by HYPERSPECTRAL V8.1 SYSTEM SD 300 and CCD-камера: B/W CAMERA BV 300 (Applied Spectral Imaging, Israel). Results interpretation was performed by GenASIs software, V8.2 (Applied Spectral Imaging, Israel).

### Chromosomal microarray analysis

Array-based comparative genomic hybridization (aCGH) was performed using SurePrint G3 Human CGH 8×60K microarrays (Agilent Technologies, Santa Clara, CA, USA) with 41 kb overall median probe spacing according to the manufacturer’s recommendations. Labelling and hybridization of the patient’s and reference DNA (#5190-3797, Human 230 Reference DNA Female, Agilent Technologies, Santa Clara, CA, USA) were performed using enzymatic labelling and hybridization protocols,v.7.5 (Agilent Technologies, Santa Clara, CA, USA). Array images were acquired with an Agilent SureScan Microarray Scanner (Agilent Technologies, Santa Clara, CA, USA). Data analysis was performed using CytoGenomics Software, v.5.1.2.1 (Agilent Technologies, Santa Clara, CA, USA) and the publicly available Database of Genomic Variants (DGV) resources. Human genome assembly 19 (hg19) was used to describe the molecular karyotype revealed by aCGH.

In addition, for some samples the CytoScan HD array (Affymetrix, USA) was applied to detect the CNVs across the entire genome following the manufacturer’s protocols. Microarray-based copy number analysis was performed using the Chromosome Analysis Suite software version 4.0 (Thermo Fisher Scientific Inc.). Detected CNVs were totally assessed by comparing them with published literature and the public databases: Database of Genomic Variants (DGV) (http://dgv.tcag.ca/dgv/app/home), DECIPHER (http://decipher.sanger.ac.uk/) and OMIM (http://www.ncbi.nlm.nih.gov/omim). Genomic positions refer to the Human Genome February 2009 assembly (GRCh37/hg19).

### Oxford Nanopore breakpoints sequencing

For nanopore sequencing, high-molecular DNA from iPS cells and blood cell samples was isolated. WGS and AS libraries were prepared using the recommended protocol for LSK109 kit from the manufacturer (Oxford Nanopore, UK). Libraries for Cas9-targeted sequencing were prepared using nCATS protocol^6^ with modifications. The sgRNAs were divided into two distinct pools similar to the “tiling” approach: the first pool of sgRNAs targeted regions spanning 10-15 kb upstream and 5-7 kb downstream from the expected breakpoints, while the second pool targeted regions located 5-7 kb upstream and 10-15 kb downstream (Supplementary Table 8), the rest of the protocol was proceeded according to nCATS method. The regions selected for AS targets included flanking areas spanning 30-40 kb from the expected breakpoints (upstream and downstream) according to the official Oxford Nanopore guidelines. Sequencing was performed on a GridION device with FLO-MIN106D flow cells (Oxford Nanopore, UK). Reads with Q-score above 9 (>90% accuracy) were taken into further analysis. Minimap2^16^ and NanoSV^7^ software were used for the alignment and variant calling processes.

### Transcriptome analysis

For transcriptome analysis, total RNA was extracted from the iPS cell differentiated to primitive streak cells and subjected to the oligo-dT-based RNA-sequencing. The resulting data were visually inspected in the IGV and analyzed using salmon^17^ and standard DESeq2 pipeline^18^.

### Exo-C data processing

All data were processed using a pipeline based on the juicer toolbox^19^, which outputs a list of pairs, hic (and cool maps made from pairs by cooler^20^, as well as quality statistics described in^3^. All analyzes were performed based on the human hg19 genome build.

### Read coverage distribution analysis

To allow comparison between coverage of Hi-C data produced by S1, DNase I, DpnII enzymes, fastq files were trimmed to 50 bp of read length and subsetted to 65 mln reads. Equalized data were aligned with BWA (v 0.7.17) (https://doi.org/10.48550/arXiv.1303.3997) on human hg38 genome build and converted to read coverage tracks with deepTools 3.5.1 bamCoverage (with the option –-binSize 1) (https://doi.org/10.1093/nar/gkw257). To construct profile plot of read density across KAPA HyperExome capture probes we excluded the segments that have DpnII sites in them and then computed coverage on ± 1 000 bp distance from center of selected capture probe segments with deepTools 3.5.1 computeMatrix reference-point (with the option –-binSize 1) and visualized it with deepTools 3.5.1 plotProfile.

To build distributions of read coverage depth exome-wide we calculated coverage sums in each capture probe segment for the same coverage tracks of the equalized data using pyBigWig 0.3.18 (https://doi.org/10.5281/zenodo.45238) and NumPy 1.21.6 (https://doi.org/10.1038/s41586-020-2649-2). The histograms of distributions were plotted using matplotlib 3.5.3 (https://doi.org/10.1109/MCSE.2007.55) and seaborn 0.13.0 (https://doi.org/10.21105/joss.03021).

## Structural variants calling

### Translocations

For translocation calling, one of two control samples was used (male control or female control). A control sample is the sum of samples of the same sex (22 female samples and 30 male samples). We combine the results of two methods for translocation calling: gradient pattern search method (works better for translocations of fragments above ∼100 kb) and line pattern search method (works better for translocations of fragments from ∼20 kb to ∼200 kb).

#### Gradient pattern search method

For each resolution of the Exo-C map (resolutions: 4096 kb, 256 kb, 16 kb, descending order), we define “dots of interest” (DOIs) as elements of Exo-C contact matrix that fulfill the following conditions: 1) DOI must have non-zero Hi-C count in both sample and control; 2) the difference of logarithms of binomial probability mass function of sample and control for the DOI is below threshold (different for each resolution);

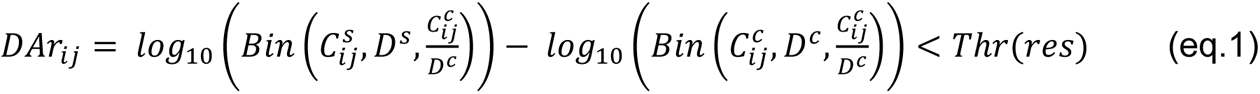

where *Bin* – binomial probability mass function, 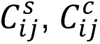 – counts at coordinates (*i*, *j*) for sample (s) and control (c); *D*^*s*^, *D*^*c*^ – sequencing depth for sample and control. To compute *D*^*s*^ and *D*^*c*^ we excluded from both control and sample Exo-C maps those dots, which contain zero value in either case or control; 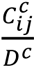– contact (*i*, *j*) probability estimation; *Thr(res)* – threshold that depends on map resolution.

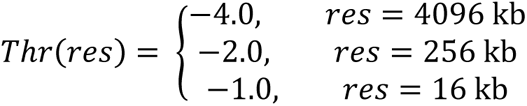

Next, for each DOI we test the hypothesis that this DOI represents contact of translocation breakpoints. Following this hypothesis, the DOI is formed by loci that are at distance 1 bin from breakpoint, their neighbors are at distance 2 bins from breakpoint, and etc. Under this assumption, each dot near DOI receives the contact probability according to the distance from the breakpoint, P_*art*_*cis*_(*m*). Under alternative assumption, there is no translocation, and dots near DOI are interchromsomal contacts with trans-contact probability P_*control*_*like*_. We estimate P_*art*_*cis*_(*m*) and P_*control*_*like*_based on dependence of contact frequencies from distance and other control sample statistics, see details in Supplementary Note 1.

Next, we compute the cumulative sum of P_*control*_*like*_ and P_*art*_*cis*_(*m*) for all dots near DOI. For DOI with coordinates *a*_0_, *b*_0_, we identify neighboring dots located within each of four rectangles anchored at DOI. We define upper right rectangle as all dots with coordinate 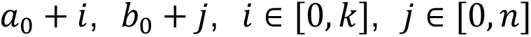; upper left rectangle as all dots with coordinate 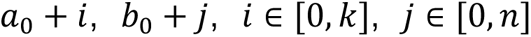; and similarly lower left and lower right rectangle. For all *k* and *n* values between 1 and 32 and each rectangle location (one of: upper right, upper left, lower right, lower left) we compute *ArLG* (Artifacts Level for Gradient pattern search) statistics (Supplementary Figure 2, E):

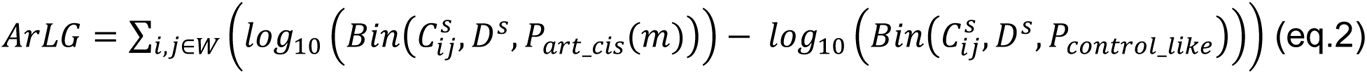

Where *W* – window, i.e. set of indices specific for the combination of rectangle location, k and n parameters; 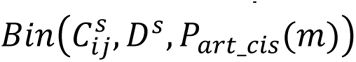 – probability for contact (*i*, *j*) to belong to the binomial distribution expected for cis-contacts of neighboring loci with distance 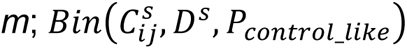 – probability for contact (*i*, *j*) to belong to the binomial distribution expected for trans-contacts; for P_*art*_*cis*_(*m*) and P_*control*_*like*_ see calculation details in Supplementary Note 1.

For each DOI, we keep only one window that gives the highest ArLG score.

After that, three filtering steps were applied:

1) *ArLG* > 0 (means that it’s more likely for contacts in a window to belong to the cis-contacts than to trans contacts);
2) If one DOI belongs to the window of another DOI, only the DOI (with its window) that has greater *ArLG* is passed;
3) log10(*ArLG)* > 0.917*log10(*contacts_sum*) – 0.92, where *contacts_sum* is sum of contacts in a window (constants were estimated from modeled translocations);

The DOIs and their windows, that passed all below steps, we considered as translocations.

#### Line pattern search method

First steps of Line pattern search method are similar to the previous one (until dots of interest are defined; resolutions: 256 kb, 16 kb, 1 kb). For each DOI, we compute the following metric

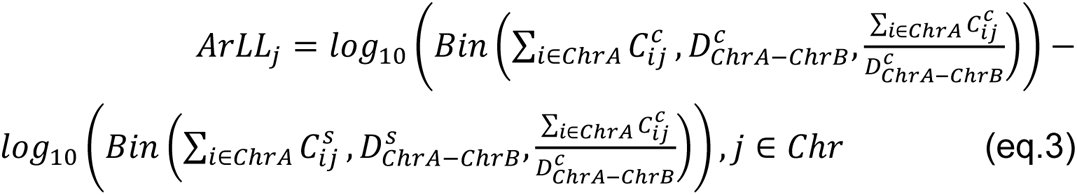

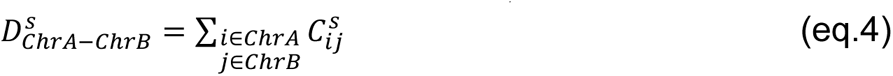

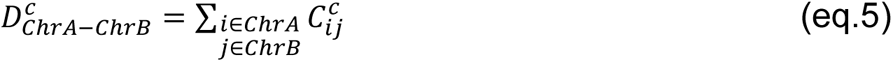

where ∑_*i*∈*C*ℎ*rA*_means the summation by Exo-C contact matrix columns that belongs to the ChrA; 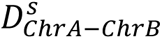 and 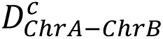 are the sum of contacts for sample and control that belong to the chromosome pair *C*ℎ*rA* − *C*ℎ*rB* (after same bins cutting, see Supplementary Note 1). For the column that is less likely to belong to sample binomial than to control binomial, the *ArLL* > 0. Every two columns were combined as one translocation when their *ArLL* > 6 and distance between them less than 5 bins. For the new column combinations the *ArLL*s were recalculated (with summation by combined columns). Additionally, the same measure was calculated by rows for every column combination to find the maximum (position that is nearest to insertion). At the end, the sample contact sum dependent filter was applied: log10(ArLL) > exp(1.73*log10(*contacts_sum*)) (constant was estimated from modeled translocations).

### Inversions

For inversion calling, we used the same control samples as for translocation calling. We called inversions at different resolutions (1 MB, 250 kb, 100 kb, 10 kb) independently and then merge all these predictions. To speed up the calculation, we have determined the minimum and maximum inversion sizes at each resolution that we are looking for (Supplementary Table 7).

For each resolution at the first stage, we computed the matrix of difference between control and experiment sample as described above in (eq. 1) formula. Further, for each dot (*i*, *j*) we computed the metric that shows how probable it is that this dot is an inversion breakpoint. This metric allows finding “butterfly”-like inversion pattern at hi-c map. We called it “sweet” metric:

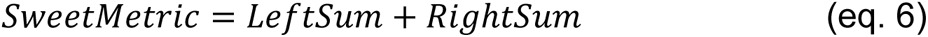

Where

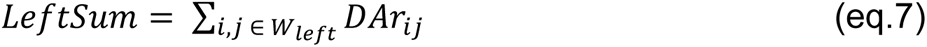

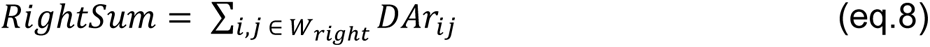

Where *DAr*_*ij*_ – the difference of logarithms of binomial probability mass function of sample and control for the dot (eq. 1). *W*_*rig*ℎ*t*_ and *W*_*left*_ – isosceles triangles with the vertex in (*i*, *j*) dot (FigSuplementSweet) and edges equal to sweet size. We used different sweet sizes for different resolutions (Table InvSizesRes).

Then we computed z-score value for each dot. We picked up the threshold for each resolution and chose all z-score values lower than this threshold. After that, we merged all overlapping coordinates at current resolution using the (*i*, *j*) coordinates of dot with the lowest z-score value as a final inversion breakpoints. Finally, we merged all predicted inversions at different resolutions and clarified each breakpoint to 10 Kb resolution. To do this we iteratively calculated “sweet” metric for dots around 10 bins from breakpoint until we reached the highest resolution 10 Kb.

### CNVs

*data preparation:* Exo-C data were aligned to the hg19 human genome assembly (https://doi.org/10.1371/journal.pbio.1001091) using BWA (v 0.7.17) (https://doi.org/10.48550/arXiv.1303.3997). Then, BAM file sorting and indexing was performed using samtools (v 1.6) (https://doi.org/10.1093/gigascience/giab008). We used DGV Gold database to exclude tool’s predictions that are possibly related to population CNV data and are not sample-specific (https://doi.org/10.1093/nar/gkt958).

*algorithms and their parameters:* We used the following tools for CNV detection based on sequence alignments: CNVkit (v 0.9.10) (https://doi.org/10.1371/journal.pcbi.1004873), CoNIFER (v 0.2.2) (http://doi.org/10.1101/gr.138115.112), GATK (v 4.4.0.0) (https://doi.org/10.1038/s41588-023-01449-0).

All tools were used with default parameters. We used 65 samples in all tool’s runs, out of which 19 or 17 (after filtration of CNVs that are less than 100,000 bp) samples formed lists of predictions to compare with control lists. Additionally, in CNVkit run one sample (P31) out of 65 was used as a normal to construct reference.

## Estimation of algorithms performance

When scoring performance of computational algorithms for SV calling, we used SV calls from validated methods (aCGH, karyotyping) as ground truth. Thus, we did not include reported SVs beyond the resolution limit of the methods (3-5 Mb for karyotyping, ∼100 kb for aCGH) in the scoring.

### EagleC performance estimation

We run EagleC^21^ with standard parameters (threshold equals to 0.9) and calculate F1-score for each type of rearrangement for 65 human samples.

In the case of translocations, we calculated all predicted rearrangements between a pair of chromosomes involved in a known translocation as one true positive value. For predicted translocations between pairs of chromosomes that do not have known rearrangement, we used the following statistics. If EagleC predicted one translocation for a pair of chromosomes, or if there were several predicted translocations between one pair of chromosomes with the distance between breakpoints more than 3 Mb, we counted each of the reported translocations as one false positive value. If the distance between breakpoints was less than 3 Mb, they were not included in assessment.

In the case of inversions, we defined true positive values as predicted rearrangements with breakpoints overlapping the 5 Mb region around known inversion breakpoints. All other predicted inversions we calculated as false positive values if the distance between breakpoints was more than 3 Mb.

Finally, in the case of CNVs, we defined true positive values as predicted rearrangements with breakpoints overlapping the 250 kb region around actual breakpoints. All other predicted CNVs we defined as false positive values if the distance between breakpoints was more than 500 kb.

### Scoring algorithm for inversion calling developed in this paper

To estimate performance of the algorithm for inversion calling, we used F1-score. We deleted repeated inversions between samples. We called all predicted inversions as true positive values if they meet following conditions:

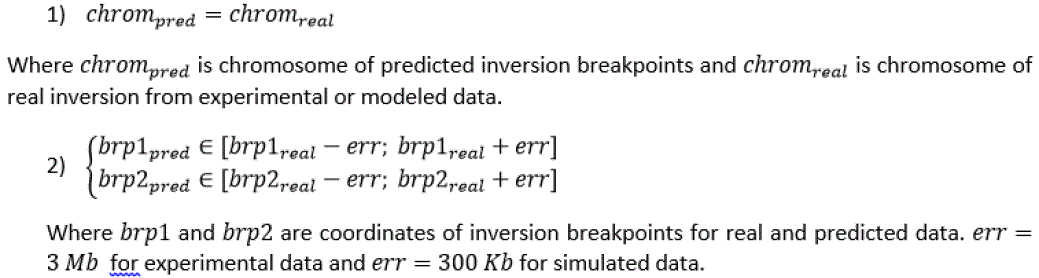

Other predicted inversions we called false positive values if their size was more than 3 Mb for experimental data and more than 20 kb for simulated data.

We defined all known inversions, which weren’t predicted, as false negative values.

### Algorithm for translocations calling

F1-score was used for estimation performance of translocation calling. Two different methods of true positives (TP) calculations were used for experimental samples and simulated samples. For experimental samples, we mark a call as TP if the chromosome pair was correct. For simulated samples, we mark a call as TP if the predicted intervals of translocation for both chromosomes intersected with the simulated translocation interval. For fragment of the chromosome *i* of the length *L* inserted into position *x* of chromosome *j* we define simulated translocation interval answer as rectangle on Exo-C contact map containing all contacts of translocated fragment with loci on chr_j with coordinates x±5*L (Supplementary Figure 2, c). If several calls were marked as TP and belong to the same chromosome pair – all of them were counted as single TP. All calls that were not marked as TP were marked as false positives (FP). If FP answers in more than 3 samples pointed at exactly the same coordinates on the same chromosome pair, they were filtered out. All translocations that were not called were marked as false negatives (FN).

### CNVkit, CoNIFER, GATK

*Preparation of known CNVs sets:* To compose CNVs control lists for tools evaluation, we classified all CNVs known to be in our samples by the source of information. This way we obtained two lists: high-confidence (clinically relevant CNVs confirmed by qPCR or other methods) and low-confidence (reported in aCGH data but not confirmed by orthogonal methods). After that we excluded repeats within lists, i.e. removed all CNVs that have intersecting boundaries. We also produced lists with filtered out CNVs whose size is less than 100,000 bp. We evaluated the quality of predictions in terms of precision and recall. We defined prediction as TP if its coordinates intersected with CNVs from control lists (allowing errors of 100,000 bp i.e. each control CNV border was extended by 100,000 bp). We also evaluated optional conditions: counting as TP only predictions which intersections with control had Jaccard Index greater than 0.5; exclusion of FP which intersections with DGV gold database had Jaccard Index greater than 0.5, or which coordinates were inside those of DGV gold database entries. Jaccard Index was calculated as: (range of intersection)/(range of union).

Precision was calculated as a proportion of all predictions marked as TP out of all predictions. To calculate recall we marked all TP, which correspond to one known CNV, as one TP, and assessed the proportion of such predictions out of known CNVs.

## Single nucleotide variants calling and annotation

SNV calling, annotation, and filtration was performed as described previously^22^. The pathogenicity of each variant was assessed according to the recommendations of the American College of Medical Genetics and Genomics (ACMG) and the Association for Molecular Pathology.

## Estimating the percentage of cells carrying a translocation in the cell mixture

To estimate the percentage of cells carrying a translocation in the in vitro cell mixtures, we used SNV data. We used GATK to call SNVs in Exo-C data for the mixture and in the NGS datasets for both types of cells present in the mixture, separately. We then identified homozygous SNVs unique to the cell type carrying a translocation using *bcftools isec* and calculated reference and alternative allele counts for each of them in the Exo-C mixture data using *samtools mpileup*. The final percentage was calculated by dividing the sum of alternative allele counts by the sum of all allele counts for each of the chosen SNV.

## Comparison of SNV recall rate between Exo-C and reference NGS methods

To evaluate Exo-C’s SNV recall rate, we compared Exo-C K562 data we generated to public reference datasets for K562, namely WGS data from^23^, Hi-C data from^24^ and^25^ and Repli-seq data from^26^. We called SNVs in all of these datasets using GATK and then filtered out the SNVs outside the exome regions using *bcftools*.

We employed the following strategy for SNV calling comparison. First, we select one of four SNV datasets as “reference”. Next, we use the remaining three datasets to construct Golden Standard SNV set. This was done by intersecting the SNV calls from the three selected datasets. We then intersected each of four versions of the Golden Standard SNVs with one remaining reference SNV dataset and with Exo-C SNV dataset. The percentage of SNVs in the intersection is shown in Supplementary Table 9. All intersections were performed using *bcftools isec*.

## Data availability

Raw human sequencing data are not available due to personalized data sharing policy. Processed Exo-C data is available under accession GEO: GSE253950.

## Code availability

Translocation and inversion calling tools developed in this study are publicly available on github: https://github.com/genomech/ExoC/.

## Acknowledgements

We acknowledge center for collective usage of computational facilities of the Institute of Cytology and Genetics (supported by budget project FWNR-2022-0019) and HPC cluster of the Novosibirsk State University (Project #2019-0546, supported by FSUS FSUS-2020-0040) for providing access to the computational resources.

Short-read sequencing of Exo-C libraries was partially supported by a grant from the Ministry of Science and Higher Education of the Russian Federation (Agreement No. 075-15-2022-310 dated April 20, 2022). Nanopore sequencing was supported by the Ministry of Science and Higher Education of the Russian Federation (Agreement 075-10-2021-093 (project GEN-RND-2017).

Chromosomes and DNA samples were obtained from “Biobank of the population of Northern Eurasia” of the Research Institute of Medical Genetics, Tomsk NRMC (http://medgenetics.ru/Biobank/). Chromosomal microarray analysis (Agilent Technology), FISH and SKY were performed on Core Facility “Medical Genomics” (Tomsk NRMC, Russia).

We thank all families for participation in the study.

