## Supplemental figures for "A novel approach for simultaneous detection of structural and single-nucleotide variants based on a combination of chromosome conformation capture and exome sequencing"

### Supplementary Figures

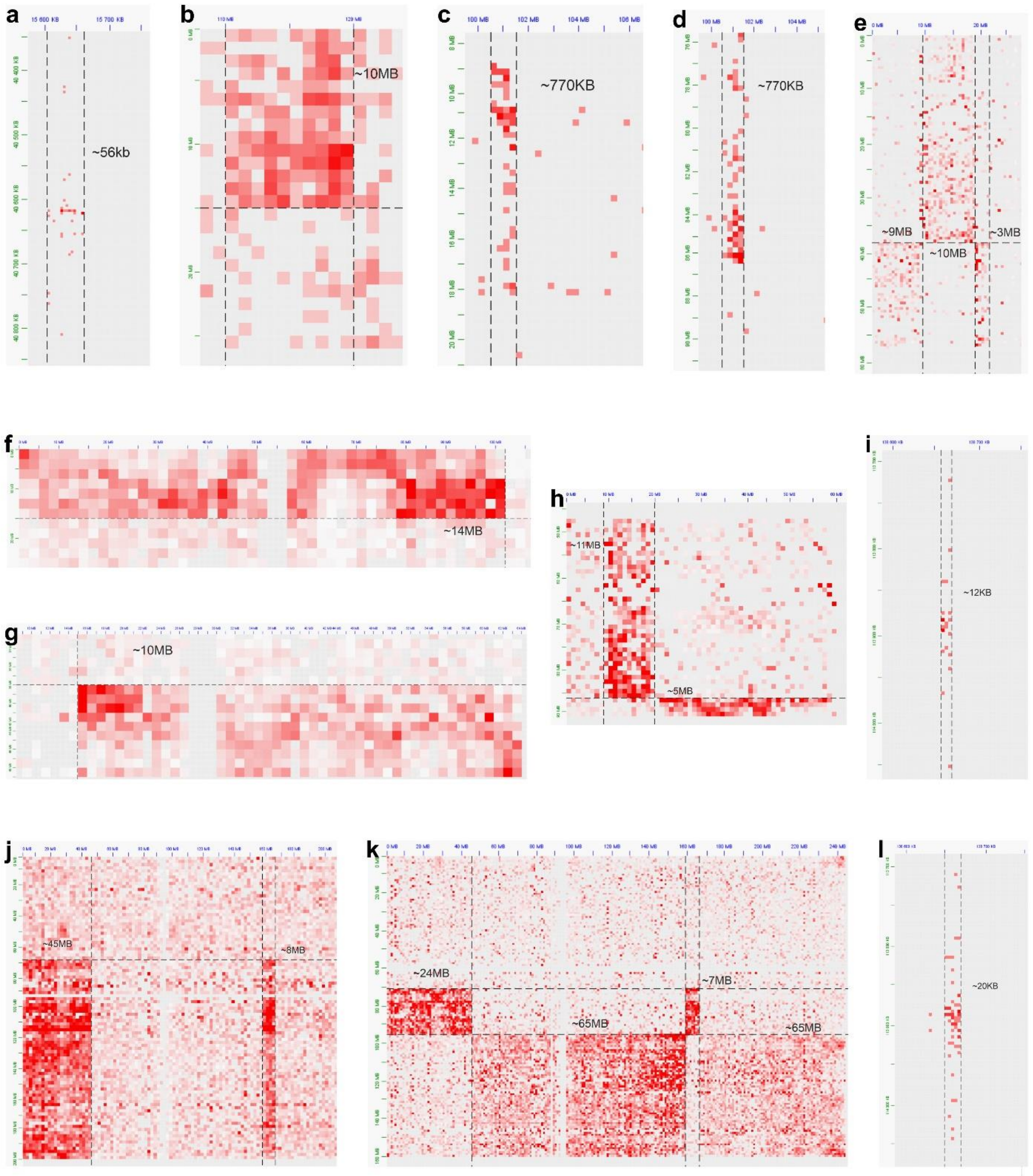


**Supplementary Figure 1**. Translocations called by Exo-C caller, but not conventional methods. Figure shows fragments of Exo-C maps for each call. Following, we provide sequence id, sample id, chromosomal pair, and resolution for each panel. **a** - s132_P10 t(4;13) 4kb; **b** - s136_P47 t(8;20)1024kb; **c** - s163_P55 t(3;6) 256kb; **d** - s163_P55 t(3;16) 256kb; **e** - s163_P55 t(6;7) 512kb; **f** - s136_P47 t(11;20) 2048kb; **g** - s136_P47 t(20;21) 1024kb; **h** - s163_P55 t(6;16) 1024kb; **i** - s210_P132 t(5;12) 4kb; **j** - s167_P63 t(2;3) 2048kb; **k** - s167_P63 t(2;7) 1024kb; **l** - s203_P119 t(5;12) 4kb.

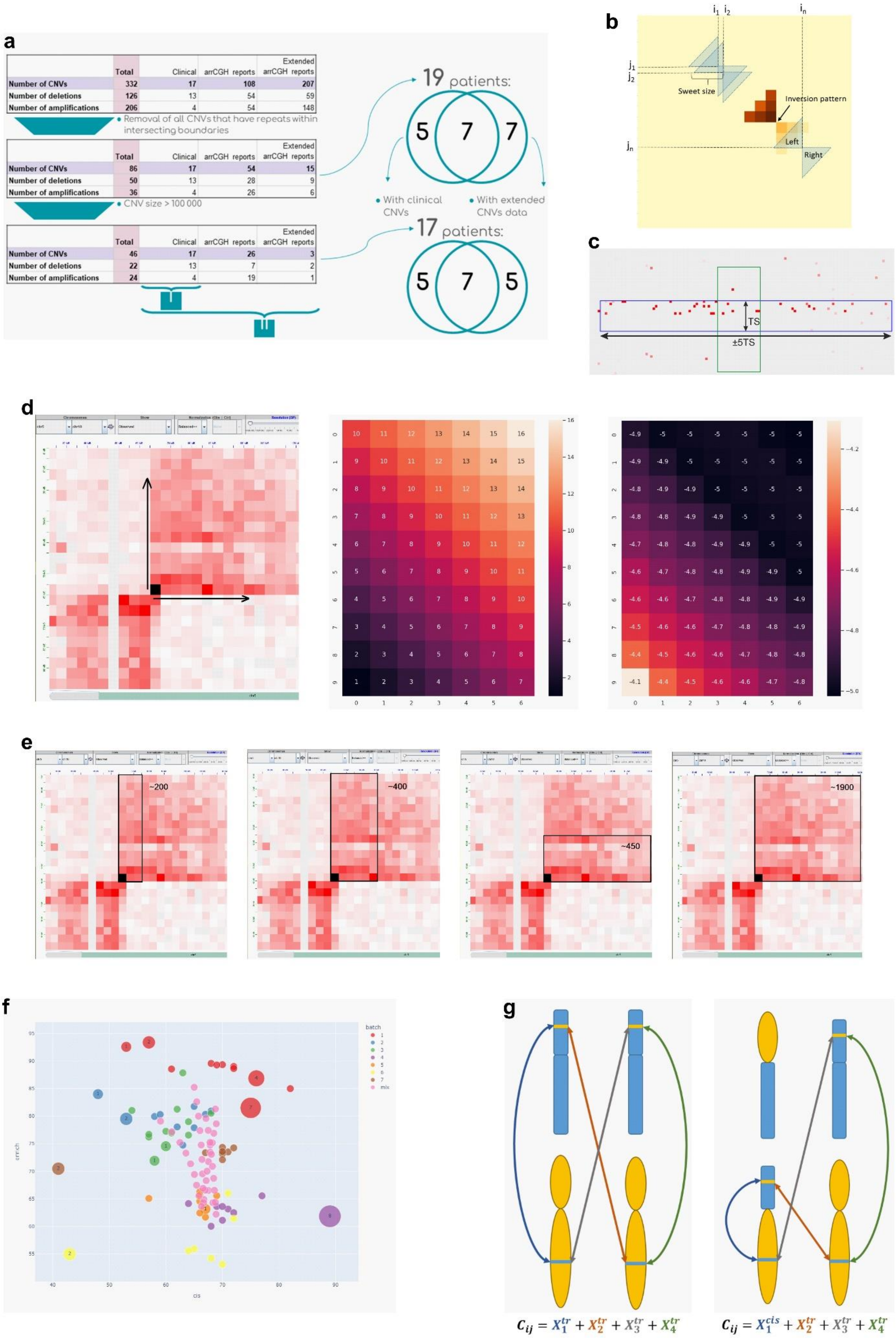
a
Total
Clinical
arrCGH reports
Extended arrCGH reports
Number of CNVs 332 17 108 207
Number of deletions 126 13 54 59
Number of amplifications 206 4 54 148
Removal of all CNVs that have repeats within intersecting boundaries
Number of CNVs 86 17 54 15
Number of deletions 50 13 28 9
Number of amplifications 36 4 26 6
CNV size > 100 000
Number of CNVs 46 17 26 3
Number of deletions 22 13 7 2
Number of amplifications 24 4 19 1
19 patients:
5 7 7
With clinical CNVs
With extended CNVs data
17 patients:
5 7 5
b
Sweet size
Inversion pattern
Left
Right
c
TS
±5TS
d
e
~200
~400
~450
~1900
f
batch
mix
g
$C_{ij} = X_1^{tr} + X_2^{tr} + X_3^{tr} + X_4^{tr}$
$C_{ij} = X_1^{cis} + X_2^{tr} + X_3^{tr} + X_4^{tr}$

**Supplementary Figure 2**. Detection of SVs using Exo-C data. **a** - Scheme, representing the process of composition and filtration of CNV lists. I — high-confidence (clinically relevant CNVs confirmed by qPCR or other methods). II — low-confidence (reported in aCGH data, but not necessarily confirmed by orthogonal methods) tiers; **b** - Schematic representation of “sweet” metric for inversion calling. (i, j) are coordinates at Hi-C matrix. It is calculating sum of values located inside of blue triangles for each dot in matrix; **c** - Example of 2D intervals intersection. Blue - simulated translocation 2D interval (width: translocation size “TS”; length: ±5 translocation sizes). Green - predicted 2D interval; **d** - Illustration of gradient pattern search method: the first pannel shows translocation pattern on Hi-C map, second pannel shows neighbors distances corresponding to corner DOI position, and third pannel shows log10(P(s)); **e** - Examples of windows and corresponding ArLG scores; **f** - The cis-enrichment distribution of samples. Each dot represents a sample. The color match the batch of experiment. The size of the dot (and the number inside dot) shows the amount of false positive translocation calls in the sample. The dots without numbers has no FPs; **g** - Schematic representation of translocation with expected number of cis- and trans-contacts showed below each scheme.

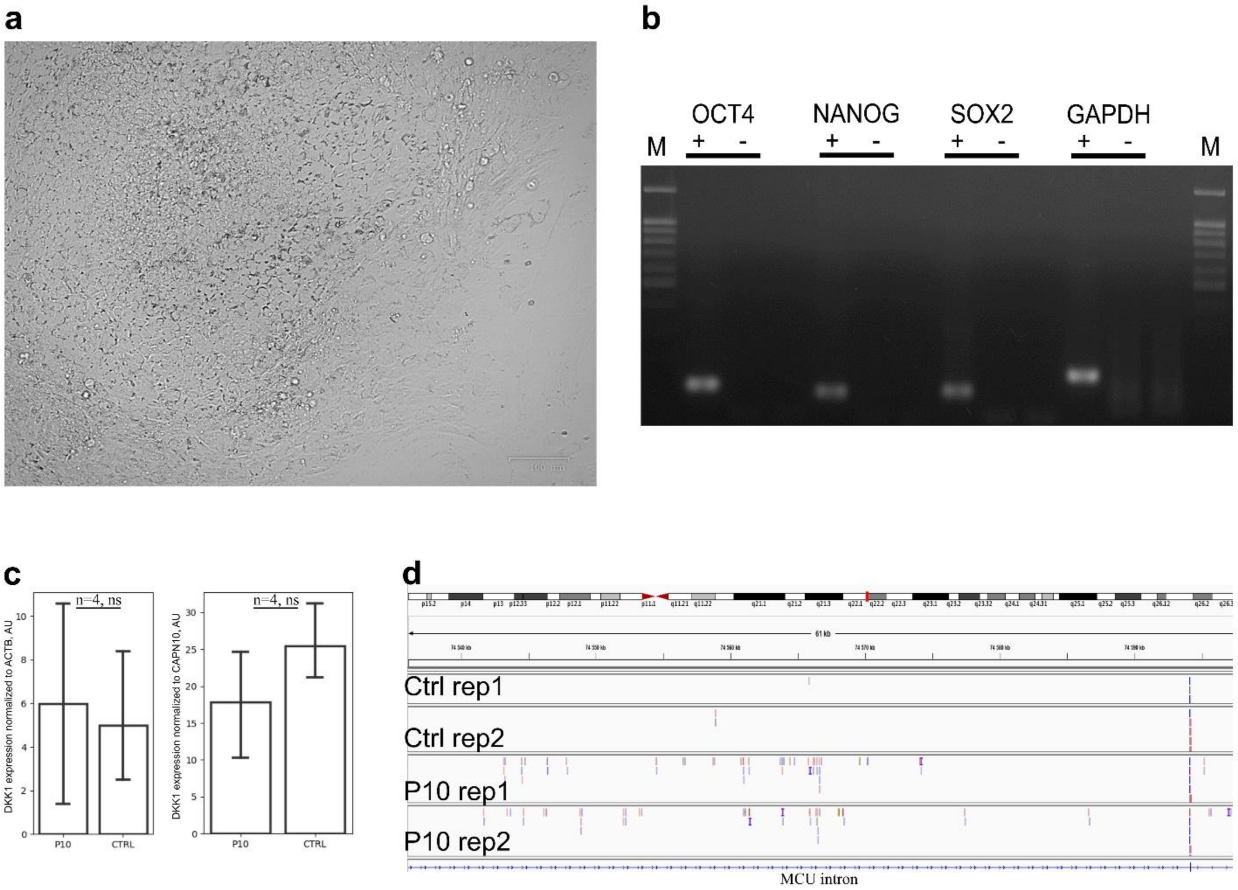


**Supplementary Figure 3**. **a** - Overview of the iPS cells obtained from P10 lymphocytes. **b** - Qualitative analysis of pluripotency gene expression in iPS cells derived from P10 lymphocytes. **c** - Digital PCR analysis of *DKK1* gene expression in primitive streak cells obtained from P10 iPS cells and control iPS cell lines. **d** - IGV screenshot showing alignment of reads obtained by transcriptome analysis of P10 iPS cells and control iPS cell lines.
