## Supplemental Tables for "A novel approach for simultaneous detection of structural and single-nucleotide variants based on a combination of chromosome conformation capture and exome sequencing"

Supplementary table 1

| Patient id | Sequence id | Sequence depth | Unmapped (%) | Duplicates (%) | HiC interactions counts | Dungling End, DE (%) | Cis contacts, (%) | Trans contacts, (%) | Capture DP > 10 | Exome Enrichment Average | Batch | Karyotype | Array CGH | WGS (Pathogenic SV calls) | Cell type | Hromatin fragmentation by |
|---|---|---|---|---|---|---|---|---|---|---|---|---|---|---|---|---|
| P26 | s90 | 44 618 032 | 0,67 | 4 | 12 354 042 | 68 | 69 | 30 | 99 | 89 | 1 | - | arr[hg19]: chr3 q13.31 (115890242_116022056)x1 | - | Lymphocytes | DNase I |
| P31 | s91 | 33 392 522 | 0,77 | 3 | 7 305 974 | 75 | 70 | 29 | 99 | 89 | 1 | - | Norm | - | Lymphocytes | DNase I |
| P32 | s92 | 31 850 565 | 0,7 | 3 | 7 737 136 | 73 | 72 | 27 | 98 | 89 | 1 | - | Norm | - | Lymphocytes | DNase I |
| P35 | s93 | 38 415 716 | 0,73 | 4 | 8 520 252 | 75 | 68 | 31 | 99 | 90 | 1 | - | arr[hg19]: chr8 p22 (15362801_15842397)×1 | - | Lymphocytes | DNase I |
| P37 | s95 | 50 365 354 | 0,85 | 4 | 10 539 758 | 76 | 53 | 46 | 99 | 93 | 1 | 46XX | arr[hg19] 8q24.11(117921970_118869338)x3 | - | Lymphocytes | DNase I |
| P38 | s96 | 51 605 033 | 0,62 | 4 | 12 728 464 | 72 | 57 | 42 | 98 | 93 | 1 | 46XX | arr[hg19] 8q24.11(117927697_118901851)x3 | - | Lymphocytes | DNase I |
| P40 | s97 | 39 255 360 | 0,76 | 4 | 10 347 472 | 70 | 72 | 27 | 98 | 89 | 1 | 46XX, inv(12)(q13.3;q24.1) | - | - | Lymphocytes | DNase I |
| P39 | s98 | 37 048 863 | 0,68 | 4 | 9 287 680 | 71 | 61 | 38 | 99 | 89 | 1 | 46XY | arr[hg19] 8p22(16798008_17889195)x1, 8p21.3p21.1(19131256_28163113)x1 | - | Lymphocytes | DNase I |
| TAF3 | s99 | 34 568 455 | 0,67 | 5 | 9 950 758 | 67 | 75 | 24 | 99 | 82 | 1 | 46XY | arr[hg19]3p26.3(1197623 −1492721)×1 | - | Fibroblasts | DNase I |
| TAF4 | s100 | 31 740 599 | 0,77 | 4 | 9 353 468 | 66 | 76 | 23 | 99 | 87 | 1 | 46XY | dup[hg19]chr3:560685-1504666 | - | Fibroblasts | DNase I |
| TAF9 | s101 | 37 631 195 | 0,74 | 4 | 9 001 544 | 73 | 82 | 17 | 99 | 85 | 1 | 46XX | [hg19](chr2:42444-2688643)x1;[hg19](chr2:2775126-24480925)x3 | - | Fibroblasts | DNase I |
| K562 | s130 | 40 684 609 | 1,2 | 18 | 12 707 020 | 58 | 81 | 18 | 98 | 85 | 2 | - | - | - | Lymphocytes | DNase I |
| P10 | s132 | 38 543 240 | 1,66 | 19 | 9 158 212 | 68 | 65 | 34 | 99 | 82 | 2 | 46XX, t(5;10) | - | - | Lymphocytes | DNase I |
| P42 | s133 | 33 723 534 | 1,45 | 18 | 8 336 228 | 67 | 48 | 51 | 99 | 84 | 2 | 46XY, t(4;12)(p14;q22)dn | - | - | Lymphocytes | DNase I |
| P43 | s134 | 32 428 792 | 1,57 | 18 | 10 500 794 | 56 | 53 | 46 | 98 | 80 | 2 | 46XY, ins(2;4) | - | - | Lymphocytes | DNase I |
| P46 | s135 | 29 902 803 | 2,11 | 16 | 9 272 774 | 59 | 58 | 41 | 99 | 80 | 2 | 46XY t(6;18;21)(p12q23; q23;q22) | - | - | Lymphocytes | DNase I |
| P47 | s136 | 28 886 122 | 1,54 | 17 | 9 270 756 | 58 | 67 | 32 | 98 | 80 | 2 | 46XY, t(8;11;21)(q23;q22;q21) | - | - | Lymphocytes | DNase I |
| P48 | s137 | 35 912 052 | 1,5 | 18 | 12 520 688 | 53 | 65 | 34 | 98 | 78 | 2 | 46XY, inv(7)(p11q11.2) | - | - | Lymphocytes | DNase I |
| P49 | s138 | 33 668 653 | 1,97 | 17 | 12 046 560 | 53 | 62 | 37 | 98 | 78 | 2 | 46XX, inv(7)(p11q11.2) | - | - | Lymphocytes | DNase I |
| P50 | s139 | 30 201 071 | 1,22 | 16 | 11 037 626 | 52 | 63 | 36 | 98 | 75 | 2 | 46XX, t(1;9)(q10.11;p10.11) | - | - | Lymphocytes | DNase I |
| P51 | s140 | 34 417 804 | 1,4 | 17 | 10 501 344 | 59 | 59 | 40 | 98 | 80 | 2 | 46XX, t(1;9)(q10.11;p10.11) | - | - | Lymphocytes | DNase I |
| P52 | s141 | 29 664 165 | 1,42 | 16 | 9 369 354 | 59 | 68 | 31 | 98 | 81 | 2 | 46XX, inv(3) | - | - | Lymphocytes | DNase I |
| P44 | s159 | 47 188 809 | 0,18 | 8 | 18 363 524 | 53 | 57 | 42 | 100 | 76 | 3 | 46XX, t(5;13)(p15;q22) | - | - | Lymphocytes | DNase I |
| P45 | s160 | 43 040 544 | 0,17 | 6 | 18 879 572 | 48 | 60 | 39 | 100 | 75 | 3 | 45XX, rob(13;14) | - | - | Lymphocytes | DNase I |
| P53 | s161 | 15 791 171 | 0,14 | 4 | 5 746 336 | 59 | 64 | 35 | 99 | 77 | 3 | Norm | - | - | Lymphocytes | DNase I |
| P54 | s162 | 43 050 094 | 0,17 | 6 | 16 001 194 | 57 | 60 | 39 | 100 | 77 | 3 | 46XY, t(2;18) | - | - | Lymphocytes | DNase I |
| P55 | s163 | 41 055 641 | 0,13 | 7 | 10 290 002 | 71 | 64 | 35 | 100 | 79 | 3 | 46XY, t(7;16)(p13;q23)dn | - | - | Lymphocytes | DNase I |
| P57 | s164 | 28 916 598 | 0,13 | 5 | 9 520 800 | 62 | 68 | 31 | 100 | 81 | 3 | 46XY, t(3;10)(p21;q11.2) | - | - | Lymphocytes | DNase I |
| P58 | s165 | 31 637 989 | 0,13 | 4 | 9 207 112 | 67 | 61 | 38 | 100 | 77 | 3 | 46XY, inv(7)(p13q21.2)pat | - | - | Lymphocytes | DNase I |
| P62 | s166 | 48 950 974 | 0,13 | 7 | 9 388 076 | 77 | 63 | 36 | 100 | 88 | 3 | 46XY, t(1;4)(p32.2;q33)dn | arr[hg19] 4q32.1q34.3(158014978_178318837)x1 | - | Lymphocytes | DNase I |
| P63 | s167 | 47 544 575 | 0,16 | 5 | 19 367 146 | 52 | 57 | 42 | 100 | 77 | 3 | 46XY, inv(2)(p21q23); t(3;7)(p13;q11.2) | - | - | Lymphocytes | DNase I |
| P69 | s168 | 38 757 892 | 0,14 | 9 | 7 620 330 | 76 | 54 | 45 | 100 | 81 | 3 | 46XY, t(2;6)(p13;q13), t(7;11)(q31.2;p15.3) | arr[hg19] 6q14.1(75917165_83621563)x1 | - | Lymphocytes | DNase I |
| P70 | s169 | 44 551 967 | 0,2 | 6 | 19 591 724 | 48 | 58 | 41 | 100 | 72 | 3 | - | - | - | Lymphocytes | DNase I |
| P71 | s170 | 51 523 861 | 0,15 | 6 | 17 179 894 | 61 | 62 | 37 | 100 | 81 | 3 | - | arr[hg19] chr5 q34(163585144_167946316)×1, chr9 p24.2p23(2267812_12275177)×1 | - | Lymphocytes | DNase I |
| P73 | s172 | 58 198 192 | 0,5 | 13 | 38 016 684 | 38 | 68 | 32 | 100 | 60 | 4 | - | - | - | Lymphocytes | S1 |
| P78 | s174 | 37 612 598 | 0,34 | 14 | 24 182 582 | 44 | 64 | 36 | 100 | 64 | 4 | 46XX, inv(8) | - | - | Lymphocytes | S1 |
| P82 | s176 | 49 774 310 | 0,6 | 18 | 31 778 607 | 47 | 70 | 30 | 100 | 64 | 4 | 46XY, inv(7) | - | - | Lymphocytes | S1 |
| P83 | s177 | 20 538 403 | 0,44 | 16 | 13 244 304 | 50 | 69 | 31 | 100 | 64 | 4 | 46XY, inv(9)(p11q12or13) | - | - | Lymphocytes | S1 |
| P84 | s178 | 43 580 686 | 0,66 | 15 | 28 623 675 | 42 | 70 | 30 | 100 | 61 | 4 | 46XY, inv(9)(p11q12or13) | - | - | Lymphocytes | S1 |
| P86 | s179 | 36 742 799 | 0,58 | 14 | 24 907 890 | 46 | 71 | 29 | 100 | 63 | 4 | 46XY, inv(9)(p11q12or13) | - | - | Lymphocytes | S1 |
| P88 | s180 | 31 352 123 | 0,35 | 15 | 21 178 727 | 48 | 72 | 28 | 99 | 62 | 4 | 46XY, inv(9)(p11q13) | - | - | Lymphocytes | S1 |
| P65 | s181 | 16 535 907 | 0,39 | 20 | 10 681 775 | 67 | 77 | 23 | 52 | 66 | 4 | 46XX | arr[hg19] Xq24-q25(118177009_123019837)×2 | - | iPSC | S1 |
| TAF16 | s182 | 63 788 738 | 0,6 | 16 | 42 221 787 | 43 | 89 | 12 | 65 | 62 | 4 | 47,XY,+mar[26]/46,XY [4].arr[GRCh38] | arr[GRCh38] 4p12p11(47818359_49218677) × 3,4q11q12(51819031_57872599) × 3 | - | Fibroblasts | S1 |
| P103 | s187 | 19 278 698 | 0,18 | 6 | 13 967 083 | 42 | 69 | 31 | 92 | 66 | 5 | 46XY, t(2;8) FISH | arr[hg19] chr2 q37.1-q37.3(232739045_243041364)×1, chr8 p23.3-p23.1(221611_919 | - | Lymphocytes | S1 |
| P104 | s188 | 21 420 552 | 0,14 | 5 | 14 886 319 | 41 | 66 | 34 | 99 | 65 | 5 | 46XY, t(2;8) FISH | - | - | Lymphocytes | S1 |
| P107 | s189 | 15 675 292 | 0,18 | 5 | 11 375 214 | 39 | 66 | 34 | 84 | 66 | 5 | 46XY, der(1)t(1;3)(q43;q26) | - | - | Lymphocytes | S1 |
| P108 | s190 | 17 177 261 | 0,17 | 5 | 12 452 744 | 36 | 66 | 34 | 91 | 62 | 5 | 46XX, t(1;3)(q43;q26) | - | - | Lymphocytes | S1 |
| P109 | s191 | 20 309 032 | 0,14 | 7 | 14 144 030 | 44 | 66 | 34 | 99 | 66 | 5 | 46XX | arr[hg19] chr1 p32.3p31.2(55199197_68901318)x1, chr10 p13p12.2(16786323_24221 | - | Lymphocytes | S1 |
| P112 | s192 | 17 508 386 | 0,17 | 5 | 12 720 868 | 42 | 57 | 43 | 94 | 65 | 5 | - | - | Norm | Lymphocytes | S1 |
| P113 | s193 | 16 594 091 | 0,16 | 5 | 11 880 080 | 36 | 67 | 34 | 92 | 63 | 5 | Norm | - | - | Lymphocytes | S1 |
| P114 | s194 | 16 560 530 | 0,15 | 4 | 12 096 851 | 35 | 67 | 33 | 97 | 62 | 5 | - | - | - | Lymphocytes | S1 |
| P91 | s197 | 15 851 276 | 0,82 | 20 | 9 701 016 | 60 | 71 | 30 | 97 | 66 | 6 | 45XX, rob(13;14)(q10;q10), inv(1)(p11,q21 | - | - | Lymphocytes | S1 |
| P92 | s198 | 19 415 322 | 0,81 | 8 | 12 938 814 | 30 | 70 | 30 | 99 | 53 | 6 | 46XX, inv(1)(p11,q21) | - | - | Lymphocytes | S1 |
| P115 | s199 | 24 615 338 | 0,86 | 8 | 16 180 674 | 32 | 68 | 32 | 99 | 54 | 6 | - | - | - | Lymphocytes | S1 |
| P119 | s203 | 60 288 598 | 0,83 | 13 | 39 682 658 | 44 | 72 | 28 | 100 | 61 | 6 | Norm | - | - | Lymphocytes | S1 |
| P122 | s204 | 26 862 328 | 0,72 | 9 | 17 371 342 | 30 | 65 | 35 | 99 | 56 | 6 | - | - | - | Lymphocytes | S1 |
| P123 | s205 | 40 893 497 | 0,71 | 9 | 25 987 926 | 33 | 64 | 36 | 100 | 56 | 6 | - | - | - | Lymphocytes | S1 |
| P124 | s206 | 27 332 293 | 0,33 | 9 | 17 409 251 | 33 | 43 | 57 | 99 | 55 | 6 | - | - | - | Lymphocytes | S1 |
| P125 | s207 | 26 356 812 | 0,35 | 11 | 18 440 696 | 40 | 67 | 33 | 100 | 73 | 7 | - | - | - | Lymphocytes | S1 |
| P131 | s209 | 23 901 997 | 0,38 | 12 | 16 815 980 | 43 | 70 | 30 | 99 | 73 | 7 | - | - | - | Lymphocytes | S1 |
| P132 | s210 | 28 586 365 | 0,33 | 13 | 19 814 877 | 41 | 71 | 29 | 100 | 74 | 7 | - | - | - | Lymphocytes | S1 |
| P133 | s211 | 20 420 473 | 0,29 | 11 | 14 408 083 | 42 | 70 | 30 | 99 | 74 | 7 | - | - | - | Lymphocytes | S1 |
| P135 | s212 | 22 696 977 | 0,31 | 11 | 16 157 617 | 42 | 70 | 30 | 99 | 74 | 7 | 46XY, t(4;9)(q34;q31) | - | - | Lymphocytes | S1 |
| P136 | s213 | 19 795 581 | 0,27 | 12 | 14 218 235 | 47 | 72 | 28 | 98 | 74 | 7 | 46XX, t(4;9)(q34;q31)pat | - | - | Lymphocytes | S1 |
| P137 | s214 | 35 272 158 | 0,36 | 15 | 23 019 540 | 38 | 41 | 59 | 100 | 70 | 7 | 46XY, inv(7)(q11.2q22) | - | - | Lymphocytes | S1 |
| P139 | s215 | 28 015 535 | 0,3 | 15 | 18 799 570 | 40 | 70 | 30 | 100 | 72 | 7 | - | - | - | Lymphocytes | S1 |

Supplementary table 2

**Found translocations**

| Patient ID | Chromosome pair | Validation result/method |
|---|---|---|
| s132_P10 | chr4-chr13 | PCR |
| s136_P47 | chr8-chr20 | Inderict confirmation by SkyFISH (chr11-chr20) and ONT (chr11-chr8 junction); see Fig. 6D |
| s136_P47 | chr11-chr20 | SKY-FISH |
| s136_P47 | chr20-chr21 | SKY-FISH |
| s163_P55 | chr3-chr6 | N/A |
| s163_P55 | chr3-chr16 | N/A |
| s163_P55 | chr6-chr7 | N/A |
| s163_P55 | chr6-chr16 | N/A |
| s167_P63 | chr2-chr3 | N/A |
| s167_P63 | chr2-chr7 | N/A |
| s203_P119 | chr5-chr12 | PCR |
| s210_P132 | chr5-chr12 | PCR |

Supplementary table 3

**Low-tier list of CNVs present in Exo-C samples**

| patient | chr | start | end | status | range |
|---|---|---|---|---|---|
| P103 | chr2 | 232739045 | 243041364 | deletion | 10302319 |
| P103 | chr8 | 221611 | 9190376 | amplification | 8968765 |
| P107 | chr11 | 5785900 | 5808616 | amplification | 22716 |
| P107 | chr1 | 247074401 | 249212668 | deletion | 2138267 |
| P107 | chr1 | 53523687 | 53675763 | deletion | 152076 |
| P107 | chr3 | 174033243 | 197840339 | amplification | 23807096 |
| P107 | chr6 | 92441400 | 96037369 | amplification | 3595969 |
| P107 | chrX | 155169996 | 155232214 | amplification | 62218 |
| P107 | chrX | 61091 | 1755786 | amplification | 1694695 |
| P109 | chr10 | 16786323 | 24221408 | deletion | 7435085 |
| P109 | chr11 | 44129525 | 44606275 | amplification | 476750 |
| P109 | chr11 | 55368154 | 55453023 | deletion | 84869 |
| P109 | chr14 | 80672550 | 80736720 | amplification | 64170 |
| P109 | chr1 | 55199197 | 68901318 | deletion | 13702121 |
| P109 | chr16 | 63711154 | 64688970 | amplification | 977816 |
| P109 | chr19 | 28407342 | 28586134 | amplification | 178792 |
| P109 | chr19 | 50086504 | 50250620 | amplification | 164116 |
| P109 | chr6 | 35203234 | 35437419 | amplification | 234185 |
| P109 | chrY | 14777314 | 23747985 | amplification | 8970671 |
| P109 | chrY | 159659 | 9940478 | amplification | 9780819 |
| P10 | chr12 | 11216816 | 11251920 | deletion | 35104 |
| P10 | chr2 | 76541402 | 76586481 | deletion | 45079 |
| P10 | chr2 | 97731119 | 98025634 | deletion | 294515 |
| P10 | chr5 | 9897966 | 9926275 | deletion | 28309 |
| P10 | chrX | 26737378 | 26823559 | deletion | 86181 |
| P10 | chrY | 6515432 | 6585532 | deletion | 70100 |
| P26 | chr3 | 115890242 | 116022056 | deletion | 131814 |
| P32 | chr12 | 31277996 | 31341928 | amplification | 63932 |
| P32 | chr19 | 35851924 | 35861544 | deletion | 9620 |
| P35 | chr8 | 15362801 | 15842397 | deletion | 479596 |

| | | | | | |
|---|---|---|---|---|---|
| P37 | chr8 | 117921970 | 118869338 | amplification | 947368 |
| P39 | chr8 | 16798008 | 17889195 | deletion | 1091187 |
| P39 | chr8 | 19131256 | 28163113 | deletion | 9031857 |
| P42 | chr1 | 148539255 | 148870387 | amplification | 331132 |
| P42 | chr11 | 5783910 | 5809230 | deletion | 25320 |
| P42 | chr1 | 196828633 | 196913251 | deletion | 84618 |
| P42 | chr12 | 7971021 | 8124048 | amplification | 153027 |
| P42 | chr16 | 16388173 | 16544419 | amplification | 156246 |
| P42 | chr1 | 72771354 | 72812440 | deletion | 41086 |
| P42 | chr6 | 67004973 | 67051189 | deletion | 46216 |
| P42 | chr7 | 34673683 | 34720673 | deletion | 46990 |
| P43 | chr11 | 49710849 | 49746642 | deletion | 35793 |
| P43 | chr14 | 35493064 | 35539847 | deletion | 46783 |
| P43 | chr17 | 43648491 | 43675217 | deletion | 26726 |
| P43 | chr5 | 97051318 | 97093707 | deletion | 42389 |
| P43 | chr7 | 38293950 | 38391352 | deletion | 97402 |
| P43 | chrX | 134753846 | 134803375 | amplification | 49529 |
| P55 | chr15 | 43889175 | 43984930 | amplification | 95755 |
| P55 | chr16 | 34466475 | 34755816 | amplification | 289341 |
| P62 | chr11 | 54969848 | 55037828 | deletion | 67980 |
| P62 | chr22 | 25656221 | 25922334 | amplification | 266113 |
| P62 | chr3 | 100340055 | 100442497 | amplification | 102442 |
| P62 | chr3 | 173239853 | 173300180 | amplification | 60327 |
| P62 | chr3 | 4183925 | 4290223 | deletion | 106298 |
| P62 | chr4 | 158014978 | 178318837 | deletion | 20303859 |
| P62 | chr4 | 81549372 | 81615070 | amplification | 65698 |
| P69 | chr12 | 86484821 | 86578187 | deletion | 93366 |
| P69 | chr16 | 32564736 | 33810309 | deletion | 1245573 |
| P69 | chr21 | 16536782 | 16939079 | deletion | 402297 |
| P69 | chr22 | 50295038 | 50326883 | deletion | 31845 |
| P69 | chr6 | 254254 | 302293 | deletion | 48039 |
| P69 | chr6 | 48337027 | 49086227 | deletion | 749200 |
| P69 | chr6 | 75917165 | 83621563 | deletion | 7704398 |
| P69 | chr7 | 54306192 | 54514155 | amplification | 207963 |

| | | | | | |
|---|---|---|---|---|---|
| P71 | chr5 | 163585144 | 167946316 | deletion | 4361172 |
| P71 | chr9 | 2267812 | 12275177 | deletion | 10007365 |
| TAF16 | chr11 | 2016675 | 2016774 | deletion | 99 |
| TAF16 | chr14 | 41616413 | 41657239 | deletion | 40826 |
| TAF16 | chr22 | 19746363 | 19754718 | amplification | 8355 |
| TAF16 | chr4 | 47820376 | 49220694 | amplification | 1400318 |
| TAF16 | chr4 | 52685197 | 58738765 | amplification | 6053568 |
| TAF3 | chr15 | 20394220 | 22669111 | amplification | 2274891 |
| TAF3 | chr20 | 1561459 | 1583221 | amplification | 21762 |
| TAF3 | chr3 | 1197623 | 1492721 | deletion | 295098 |
| TAF4 | chr11 | 18951965 | 18963642 | deletion | 11677 |
| TAF4 | chr1 | 196744721 | 196804417 | deletion | 59696 |
| TAF4 | chr14 | 106331956 | 107216227 | deletion | 884271 |
| TAF4 | chr14 | 75106700 | 75116568 | deletion | 9868 |
| TAF4 | chr2 | 89163862 | 89319978 | deletion | 156116 |
| TAF4 | chr3 | 560685 | 1504666 | amplification | 943981 |
| TAF4 | chr7 | 26888579 | 26932834 | deletion | 44255 |
| TAF9 | chr12 | 31277996 | 31341928 | deletion | 63932 |
| TAF9 | chr19 | 35851924 | 35861544 | amplification | 9620 |
| TAF9 | chr2 | 2775126 | 24480925 | amplification | 21705799 |
| TAF9 | chr2 | 42444 | 2688643 | deletion | 2646199 |
| TAF9 | chr3 | 4064411 | 4153829 | amplification | 89418 |

Supplementary table 4

**Results of CNV prediction tools evaluation with different conditions.**

| Tool | Precision | Recall | TP+FP | TP | TPunique | ControlList | SizeFiltration | RelevantItemsNumber | PredictionControlIntersectionFiltration | dgvGoldExcludedFromFP |
|---|---|---|---|---|---|---|---|---|---|---|
| GATK | 0.013893129770992366 | 0.36046511627906974 | 6550 | 91 | 31 | Low-tier | No | 86 | No | No |
| GATK | 0.20973782771535582 | 0.3695652173913043 | 267 | 56 | 17 | Low-tier | > 100000 bp | 46 | No | No |
| GATK | 0.0006106870229007634 | 0.046511627906976744 | 6550 | 4 | 4 | Low-tier | No | 86 | Jaccard Index > 0.5 | No |
| GATK | 0.0149812734082397 | 0.08695652173913043 | 267 | 4 | 4 | Low-tier | > 100000 bp | 46 | Jaccard Index > 0.5 | No |
| GATK | 0.0019120458891013384 | 0.046511627906976744 | 2092 | 4 | 4 | Low-tier | No | 86 | Jaccard Index > 0.5 | Yes |
| GATK | 0.016597510373443983 | 0.08695652173913043 | 241 | 4 | 4 | Low-tier | > 100000 bp | 46 | Jaccard Index > 0.5 | Yes |
| GATK | 0.011060507482108002 | 0.7058823529411765 | 4611 | 51 | 12 | High-tier | No | 17 | No | No |
| GATK | 0.2388888888888889 | 0.6470588235294118 | 180 | 43 | 11 | High-tier | > 100000 bp | 17 | No | No |
| GATK | 0.00043374539145521576 | 0.11764705882352941 | 4611 | 2 | 2 | High-tier | No | 17 | Jaccard Index > 0.5 | No |
| GATK | 0.011111111111111112 | 0.11764705882352941 | 180 | 2 | 2 | High-tier | > 100000 bp | 17 | Jaccard Index > 0.5 | No |
| GATK | 0.0014194464158978 | 0.11764705882352941 | 1409 | 2 | 2 | High-tier | No | 17 | Jaccard Index > 0.5 | Yes |
| GATK | 0.013513513513513514 | 0.11764705882352941 | 148 | 2 | 2 | High-tier | > 100000 bp | 17 | Jaccard Index > 0.5 | Yes |
| CoNIFER | 0.06500607533414338 | 0.2441860465116279 | 1646 | 107 | 21 | Low-tier | No | 86 | No | No |
| CoNIFER | 0.20161290322580644 | 0.391304347826087 | 372 | 75 | 18 | Low-tier | > 100000 bp | 46 | No | No |
| CoNIFER | 0.00425273390036452 | 0.08139534883720931 | 1646 | 7 | 7 | Low-tier | No | 86 | Jaccard Index > 0.5 | No |
| CoNIFER | 0.01881720430107527 | 0.15217391304347827 | 372 | 7 | 7 | Low-tier | > 100000 bp | 46 | Jaccard Index > 0.5 | No |
| CoNIFER | 0.005216095380029807 | 0.08139534883720931 | 1342 | 7 | 7 | Low-tier | No | 86 | Jaccard Index > 0.5 | Yes |
| CoNIFER | 0.020588235294117647 | 0.15217391304347827 | 340 | 7 | 7 | Low-tier | > 100000 bp | 46 | Jaccard Index > 0.5 | Yes |
| CoNIFER | 0.053353658536585365 | 0.7058823529411765 | 1312 | 70 | 12 | High-tier | No | 17 | No | No |
| CoNIFER | 0.16666666666666666 | 0.7058823529411765 | 330 | 55 | 12 | High-tier | > 100000 bp | 17 | No | No |
| CoNIFER | 0.003048780487804878 | 0.23529411764705882 | 1312 | 4 | 4 | High-tier | No | 17 | Jaccard Index > 0.5 | No |
| CoNIFER | 0.012121212121212121 | 0.23529411764705882 | 330 | 4 | 4 | High-tier | > 100000 bp | 17 | Jaccard Index > 0.5 | No |
| CoNIFER | 0.0037105751391465678 | 0.23529411764705882 | 1078 | 4 | 4 | High-tier | No | 17 | Jaccard Index > 0.5 | Yes |
| CoNIFER | 0.013513513513513514 | 0.23529411764705882 | 296 | 4 | 4 | High-tier | > 100000 bp | 17 | Jaccard Index > 0.5 | Yes |
| CNVkit | 0.014992503748125937 | 0.22093023255813954 | 1334 | 20 | 19 | Low-tier | No | 86 | No | No |
| CNVkit | 0.01778242677824268 | 0.34782608695652173 | 956 | 17 | 16 | Low-tier | > 100000 bp | 46 | No | No |
| CNVkit | 0.0074962518740629685 | 0.11627906976744186 | 1334 | 10 | 10 | Low-tier | No | 86 | Jaccard Index > 0.5 | No |
| CNVkit | 0.010460251046025104 | 0.21739130434782608 | 956 | 10 | 10 | Low-tier | > 100000 bp | 46 | Jaccard Index > 0.5 | No |
| CNVkit | 0.008460236886632826 | 0.11627906976744186 | 1182 | 10 | 10 | Low-tier | No | 86 | Jaccard Index > 0.5 | Yes |
| CNVkit | 0.009823182711198428 | 0.21739130434782608 | 1018 | 10 | 10 | Low-tier | > 100000 bp | 46 | Jaccard Index > 0.5 | Yes |
| CNVkit | 0.010648596321393998 | 0.5882352941176471 | 1033 | 11 | 10 | High-tier | No | 17 | No | No |
| CNVkit | 0.012835472578763127 | 0.5882352941176471 | 857 | 11 | 10 | High-tier | > 100000 bp | 17 | No | No |
| CNVkit | 0.006776379477250726 | 0.4117647058823529 | 1033 | 7 | 7 | High-tier | No | 17 | Jaccard Index > 0.5 | No |
| CNVkit | 0.008168028004667444 | 0.4117647058823529 | 857 | 7 | 7 | High-tier | > 100000 bp | 17 | Jaccard Index > 0.5 | No |
| CNVkit | 0.007543103448275862 | 0.4117647058823529 | 928 | 7 | 7 | High-tier | No | 17 | Jaccard Index > 0.5 | Yes |
| CNVkit | 0.008739076154806492 | 0.4117647058823529 | 801 | 7 | 7 | High-tier | > 100000 bp | 17 | Jaccard Index > 0.5 | Yes |

Supplementary table 5

**SamplesMix**

| Sample with normal karyotype | Sample with SV | Chromosomal pairs with translocation | Cells/data with SV (percentage) | Experiment type |
|---|---|---|---|---|
| P73 | P10 | chr5-chr10, chr4-chr13 | 10% - 90% | *in silico* |
| P113 | P47 | chr8-chr11, chr8-chr21, chr8-chr20, chr11-chr20, chr20-chr21 | 10% - 90% | *in silico* |
| P114 | P62 | chr1-chr4 | 10% - 90% | *in silico* |
| P114 | P69 | chr2-chr6, chr7-chr11 | 10% - 90% | *in silico* |
| P114 | P62 | chr1-chr4 | 10%, 50%, 75% | *In vitro* |
| P114 | P69 | chr2-chr6, chr7-chr11 | 25% | *In vitro* |

Supplementary table 6

### Validation of breakpoints found via Exo-C with nanopore sequencing.

| Sample | Breakpoints detected with | Previously validated | Validated with Nanopore by manual analysis of split-reads | | | | Total yield, Gb (Q-score > 9) | | |
|---|---|---|---|---|---|---|---|---|---|
| | | | Total | WGS | AS | nCATS | WGS | AS | nCATS |
| P10 | 4 | 2 (PCR/FISH) | 4 | 1 (0) | 3 (1) | - | 45048 | 1.32 | - |
| P43 | 7 | 6 (FISH) | 7 | 1 (0) | 5 (0) | 5 (4) | 4 | 0.57 | 45109 |
| P47 | 6 | 3 (FISH) | 3 | 1 (0) | 2 (1) | - | 2.64 | 1.18 | - |
| P63 | 4 | 4 (Karyotyping) | 2 | 1 (0) | 2 (0) | 2 (0) | 1.96 | 1.25 | 2.52 |
| P82 | 2 | 2 (Karyotyping) | 2 | 1 (0) | 2 (0) | - | 3.45 | 0.42 | - |

Supplementary table 7

**Sizes of called inversions at different resolutions (bin sizes)**

| Resolution (bp) | Minimum inversion size (bp) | Maximum inversion size (bp) | Sweet size (bins) |
|---|---|---|---|
| 1000000 | 25000000 | Chromosome size | 5 |
| 250000 | 10000000 | 30000000 | 10 |
| 100000 | 2000000 | 12000000 | 10 |
| 10000 | 50000 | 3000000 | 20 |

Supplementary table 8

**List of primers used for sgRNA generation**

| Primer | Sequence (5' – 3') | Description |
|---|---|---|
| SpCas9-scaffold-R | aaaagcaccgactcggtgcc | Reverse primer for amplification of sgRNA template from scaffold sequence in PX458 plasmid (Addgene #48138) |
| P43_A1-1 | aagcTAATACGACTCACTATAGGTATTCCTCACAATTCAAAGTGTTTTAGAGCTAGAAATAGCAAGTTAA | P43 case. Forward primer for amplification of sgRNA template. Includes T7 promoter sequence for in vitro transcription. |
| P43_A1-2 | aagcTAATACGACTCACTATAGGCTGAGTGTCGTACTCACCTCGTTTTAGAGCTAGAAATAGCAAGTTAA | P43 case. Forward primer for amplification of sgRNA template. Includes T7 promoter sequence for in vitro transcription. |
| P43_A2-1 | aagcTAATACGACTCACTATAGGCAGTCGAATTCTAAGCAATTGTTTTAGAGCTAGAAATAGCAAGTTAA | P43 case. Forward primer for amplification of sgRNA template. Includes T7 promoter sequence for in vitro transcription. |
| P43_A2-2 | aagcTAATACGACTCACTATAGGGTATTCATACCTAACATGGTTTTAGAGCTAGAAATAGCAAGTTAA | P43 case. Forward primer for amplification of sgRNA template. Includes T7 promoter sequence for in vitro transcription. |
| P43_B1-1 | aagcTAATACGACTCACTATAGGTGTGACCTCTGATACTGAGCGTTTTAGAGCTAGAAATAGCAAGTTAA | P43 case. Forward primer for amplification of sgRNA template. Includes T7 promoter sequence for in vitro transcription. |
| P43_B1-2 | aagcTAATACGACTCACTATAGGTTTGAACTCCTGCATGAACAGTTTTAGAGCTAGAAATAGCAAGTTAA | P43 case. Forward primer for amplification of sgRNA template. Includes T7 promoter sequence for in vitro transcription. |
| P43_B2-1 | aagcTAATACGACTCACTATAGGCCTTACTTAGGGTAGTGCCAGTTTTAGAGCTAGAAATAGCAAGTTAA | P43 case. Forward primer for amplification of sgRNA template. Includes T7 promoter sequence for in vitro transcription. |
| P43_B2-2 | aagcTAATACGACTCACTATAGGTGGCCAAAAATAGTGAAGTCGTTTTAGAGCTAGAAATAGCAAGTTAA | P43 case. Forward primer for amplification of sgRNA template. Includes T7 promoter sequence for in vitro transcription. |
| P43_C1-1 | aagcTAATACGACTCACTATAGGCTTAGGTGTCTACTTCCCTTGTTTTAGAGCTAGAAATAGCAAGTTAA | P43 case. Forward primer for amplification of sgRNA template. Includes T7 promoter sequence for in vitro transcription. |
| P43_C1-2 | aagcTAATACGACTCACTATAGGTTGTTTCTATACAATATGGCGTTTTAGAGCTAGAAATAGCAAGTTAA | P43 case. Forward primer for amplification of sgRNA template. Includes T7 promoter sequence for in vitro transcription. |
| P43_C1-3 | aagcTAATACGACTCACTATAGGTGACACGCCATGTTTCCTATGTTTTAGAGCTAGAAATAGCAAGTTAA | P43 case. Forward primer for amplification of sgRNA template. Includes T7 promoter sequence for in vitro transcription. |
| P43_C1-4 | aagcTAATACGACTCACTATAGGTGACCACCAAACTGGCAGCTGTTTTAGAGCTAGAAATAGCAAGTTAA | P43 case. Forward primer for amplification of sgRNA template. Includes T7 promoter sequence for in vitro transcription. |
| P43_C2-1 | aagcTAATACGACTCACTATAGGTAGACCTGTCTCTACCACTAGTTTTAGAGCTAGAAATAGCAAGTTAA | P43 case. Forward primer for amplification of sgRNA template. Includes T7 promoter sequence for in vitro transcription. |
| P43_C2-2 | aagcTAATACGACTCACTATAGGACACGTAGGATTCATATTGAGTTTTAGAGCTAGAAATAGCAAGTTAA | P43 case. Forward primer for amplification of sgRNA template. Includes T7 promoter sequence for in vitro transcription. |
| P43_C2-3 | aagcTAATACGACTCACTATAGGAGGCGAGTCTTATGCTAACTGTTTTAGAGCTAGAAATAGCAAGTTAA | P43 case. Forward primer for amplification of sgRNA template. Includes T7 promoter sequence for in vitro transcription. |
| P43_C2-4 | aagcTAATACGACTCACTATAGGAACTGAGTCTTGTACTAACCGTTTTAGAGCTAGAAATAGCAAGTTAA | P43 case. Forward primer for amplification of sgRNA template. Includes T7 promoter sequence for in vitro transcription. |
| P43_D1-1 | aagcTAATACGACTCACTATAGGTACGTATCATTCTTTTTTGTTTTAGAGCTAGAAATAGCAAGTTAA | P43 case. Forward primer for amplification of sgRNA template. Includes T7 promoter sequence for in vitro transcription. |
| P43_D1-2 | aagcTAATACGACTCACTATAGGTGGATTGTATAGAGCATACGTTTTAGAGCTAGAAATAGCAAGTTAA | P43 case. Forward primer for amplification of sgRNA template. Includes T7 promoter sequence for in vitro transcription. |

| | | |
|---|---|---|
| P43_D2-1 | aagcTAATACGACTCACTATAGGTACTGAATAAGATCGAACAGTTTTAGAGCTAGAAATAGCAAGTTAA | P43 case. Forward primer for amplification of sgRNA template. Includes T7 promoter sequence for in vitro transcription. |
| P43_D2-2 | aagcTAATACGACTCACTATAGGAGTCTTCTATCAAATTCAGGGTTTTAGAGCTAGAAATAGCAAGTTAA | P43 case. Forward primer for amplification of sgRNA template. Includes T7 promoter sequence for in vitro transcription. |
| P43_E1-1 | aagcTAATACGACTCACTATAGGAGTTGATCATATATGATACTGTTTTAGAGCTAGAAATAGCAAGTTAA | P43 case. Forward primer for amplification of sgRNA template. Includes T7 promoter sequence for in vitro transcription. |
| P43_E1-2 | aagcTAATACGACTCACTATAGGCAGTGAATTATGGTACTCTCGTTTTAGAGCTAGAAATAGCAAGTTAA | P43 case. Forward primer for amplification of sgRNA template. Includes T7 promoter sequence for in vitro transcription. |
| P43_E1-3 | aagcTAATACGACTCACTATAGGCGTTGTCCTTCTCCATACATGTTTTAGAGCTAGAAATAGCAAGTTAA | P43 case. Forward primer for amplification of sgRNA template. Includes T7 promoter sequence for in vitro transcription. |
| P43_E1-4 | aagcTAATACGACTCACTATAGGTTGGGCTCTGCATCCCCGAGGTTTTAGAGCTAGAAATAGCAAGTTAA | P43 case. Forward primer for amplification of sgRNA template. Includes T7 promoter sequence for in vitro transcription. |
| P43_E2-1 | aagcTAATACGACTCACTATAGGATGGGTAGGATATAAGATCGTTTTAGAGCTAGAAATAGCAAGTTAA | P43 case. Forward primer for amplification of sgRNA template. Includes T7 promoter sequence for in vitro transcription. |
| P43_E2-2 | aagcTAATACGACTCACTATAGGACTAACTATTACAGATCACAGTTTTAGAGCTAGAAATAGCAAGTTAA | P43 case. Forward primer for amplification of sgRNA template. Includes T7 promoter sequence for in vitro transcription. |
| P43_E2-3 | aagcTAATACGACTCACTATAGGACCTCCTATAAAGAAACGATGTTTTAGAGCTAGAAATAGCAAGTTAA | P43 case. Forward primer for amplification of sgRNA template. Includes T7 promoter sequence for in vitro transcription. |
| P43_E2-4 | aagcTAATACGACTCACTATAGGAATGAATTGAGTTCCTTCTGTTTTAGAGCTAGAAATAGCAAGTTAA | P43 case. Forward primer for amplification of sgRNA template. Includes T7 promoter sequence for in vitro transcription. |
| P43_F1-1 | aagcTAATACGACTCACTATAGGATAGACCCTAATTGACCCCAGTTTTAGAGCTAGAAATAGCAAGTTAA | P43 case. Forward primer for amplification of sgRNA template. Includes T7 promoter sequence for in vitro transcription. |
| P43_F1-2 | aagcTAATACGACTCACTATAGGCTCATTGATAGTCCCGAGTGGTTTTAGAGCTAGAAATAGCAAGTTAA | P43 case. Forward primer for amplification of sgRNA template. Includes T7 promoter sequence for in vitro transcription. |
| P43_F2-1 | aagcTAATACGACTCACTATAGGGTGGTGAACTATCATCAAGTTTTAGAGCTAGAAATAGCAAGTTAA | P43 case. Forward primer for amplification of sgRNA template. Includes T7 promoter sequence for in vitro transcription. |
| P43_F2-2 | aagcTAATACGACTCACTATAGGAAGGGAGCAGGATACTTTCGTTTTAGAGCTAGAAATAGCAAGTTAA | P43 case. Forward primer for amplification of sgRNA template. Includes T7 promoter sequence for in vitro transcription. |
| P43_G1-1 | aagcTAATACGACTCACTATAGGATCTACCATATTACAAGCTAGTTTTAGAGCTAGAAATAGCAAGTTAA | P43 case. Forward primer for amplification of sgRNA template. Includes T7 promoter sequence for in vitro transcription. |
| P43_G1-2 | aagcTAATACGACTCACTATAGGTTCTGACGACACGATTCCGTTTTAGAGCTAGAAATAGCAAGTTAA | P43 case. Forward primer for amplification of sgRNA template. Includes T7 promoter sequence for in vitro transcription. |
| P43_G2-1 | aagcTAATACGACTCACTATAGGCAATACCTAAGTTGTCATGTTTTAGAGCTAGAAATAGCAAGTTAA | P43 case. Forward primer for amplification of sgRNA template. Includes T7 promoter sequence for in vitro transcription. |
| P43_G2-2 | aagcTAATACGACTCACTATAGGTATAAGTCGTACCTTCCTATGTTTTAGAGCTAGAAATAGCAAGTTAA | P43 case. Forward primer for amplification of sgRNA template. Includes T7 promoter sequence for in vitro transcription. |
| P63_A1-1 | aagcTAATACGACTCACTATAGGCGCAAGGTAGAGAGTATTGTTTTAGAGCTAGAAATAGCAAGTTAA | P63 case. Forward primer for amplification of sgRNA template. Includes T7 promoter sequence for in vitro transcription. |
| P63_A1-2 | aagcTAATACGACTCACTATAGGTAGGCCCATTTTGTTGAGAGGTTTTAGAGCTAGAAATAGCAAGTTAA | P63 case. Forward primer for amplification of sgRNA template. Includes T7 promoter sequence for in vitro transcription. |
| P63_A2-1 | aagcTAATACGACTCACTATAGGCGCTTTAGTAAATGATGAATGTTTTAGAGCTAGAAATAGCAAGTTAA | P63 case. Forward primer for amplification of sgRNA template. Includes T7 promoter sequence for in vitro transcription. |

| | | |
|---|---|---|
| P63_A2-2 | aagcTAATACGACTCACTATAGGCTTGTTTATGTACCTCGTTTGTTTTAGAGCTAGAAATAGCAAGTTAA | P63 case. Forward primer for amplification of sgRNA template. Includes T7 promoter sequence for in vitro transcription. |
| P63_A3-1 | aagcTAATACGACTCACTATAGGAGATGTTGACATGTGTGCCGGTTTTAGAGCTAGAAATAGCAAGTTAA | P63 case. Forward primer for amplification of sgRNA template. Includes T7 promoter sequence for in vitro transcription. |
| P63_A3-2 | aagcTAATACGACTCACTATAGGTCAGGGCTGTAAATTAGACGTTTTAGAGCTAGAAATAGCAAGTTAA | P63 case. Forward primer for amplification of sgRNA template. Includes T7 promoter sequence for in vitro transcription. |
| P63_B1-1 | aagcTAATACGACTCACTATAGGTTGCAAGAACTAATAACTCCGTTTTAGAGCTAGAAATAGCAAGTTAA | P63 case. Forward primer for amplification of sgRNA template. Includes T7 promoter sequence for in vitro transcription. |
| P63_B1-2 | aagcTAATACGACTCACTATAGGGCGTAGGTTTTACACTTAGTTTTAGAGCTAGAAATAGCAAGTTAA | P63 case. Forward primer for amplification of sgRNA template. Includes T7 promoter sequence for in vitro transcription. |
| P63_B2-1 | aagcTAATACGACTCACTATAGGTATGTCAACCTTGTAGAGACGTTTTAGAGCTAGAAATAGCAAGTTAA | P63 case. Forward primer for amplification of sgRNA template. Includes T7 promoter sequence for in vitro transcription. |
| P63_B2-2 | aagcTAATACGACTCACTATAGGTGTCTACTTCATATGGGGTTGTTTTAGAGCTAGAAATAGCAAGTTAA | P63 case. Forward primer for amplification of sgRNA template. Includes T7 promoter sequence for in vitro transcription. |
| P63_B3-1 | aagcTAATACGACTCACTATAGGCAATCCTACAAGTACCCAGCGTTTTAGAGCTAGAAATAGCAAGTTAA | P63 case. Forward primer for amplification of sgRNA template. Includes T7 promoter sequence for in vitro transcription. |
| P63_B3-2 | aagcTAATACGACTCACTATAGGCTCCATTCTGGATCAAACGAGTTTTAGAGCTAGAAATAGCAAGTTAA | P63 case. Forward primer for amplification of sgRNA template. Includes T7 promoter sequence for in vitro transcription. |
| P63_C1-1 | aagcTAATACGACTCACTATAGGCTATCTTTAGTGCATATCCGTTTTAGAGCTAGAAATAGCAAGTTAA | P63 case. Forward primer for amplification of sgRNA template. Includes T7 promoter sequence for in vitro transcription. |
| P63_C1-2 | aagcTAATACGACTCACTATAGGCATGGCTGCAACCAACCGGGGTTTTAGAGCTAGAAATAGCAAGTTAA | P63 case. Forward primer for amplification of sgRNA template. Includes T7 promoter sequence for in vitro transcription. |
| P63_C2-1 | aagcTAATACGACTCACTATAGGTATGGGTCAAAACACAGTCTGTTTTAGAGCTAGAAATAGCAAGTTAA | P63 case. Forward primer for amplification of sgRNA template. Includes T7 promoter sequence for in vitro transcription. |
| P63_C2-2 | aagcTAATACGACTCACTATAGGCACTCGTCACTTAGTTTTGGGTTTTAGAGCTAGAAATAGCAAGTTAA | P63 case. Forward primer for amplification of sgRNA template. Includes T7 promoter sequence for in vitro transcription. |
| P63_C3-1 | aagcTAATACGACTCACTATAGGTCATCTGTTCGTTCCGATCAGTTTTAGAGCTAGAAATAGCAAGTTAA | P63 case. Forward primer for amplification of sgRNA template. Includes T7 promoter sequence for in vitro transcription. |
| P63_C3-2 | aagcTAATACGACTCACTATAGGCCCCAATAAATGCTATGCAGGGTTTTAGAGCTAGAAATAGCAAGTTAA | P63 case. Forward primer for amplification of sgRNA template. Includes T7 promoter sequence for in vitro transcription. |
| P63_D1-1 | aagcTAATACGACTCACTATAGGCGTAATCTCAGGAATTCCAGTTTTAGAGCTAGAAATAGCAAGTTAA | P63 case. Forward primer for amplification of sgRNA template. Includes T7 promoter sequence for in vitro transcription. |
| P63_D1-2 | aagcTAATACGACTCACTATAGGCACGGATGGATTTCGGTGCAGTTTTAGAGCTAGAAATAGCAAGTTAA | P63 case. Forward primer for amplification of sgRNA template. Includes T7 promoter sequence for in vitro transcription. |
| P63_D2-1 | aagcTAATACGACTCACTATAGGCAGTAGGCATATTCTCTGTGTTTTAGAGCTAGAAATAGCAAGTTAA | P63 case. Forward primer for amplification of sgRNA template. Includes T7 promoter sequence for in vitro transcription. |
| P63_D2-2 | aagcTAATACGACTCACTATAGGTCTAAGCACAACTTTTTGGGTTTTAGAGCTAGAAATAGCAAGTTAA | P63 case. Forward primer for amplification of sgRNA template. Includes T7 promoter sequence for in vitro transcription. |
| P63_D3-1 | aagcTAATACGACTCACTATAGGTTAGGCTGTTAGTGTGAACGTTTTAGAGCTAGAAATAGCAAGTTAA | P63 case. Forward primer for amplification of sgRNA template. Includes T7 promoter sequence for in vitro transcription. |
| P63_D3-2 | aagcTAATACGACTCACTATAGGATATCCACGCGCTGACAAGTTTTAGAGCTAGAAATAGCAAGTTAA | P63 case. Forward primer for amplification of sgRNA template. Includes T7 promoter sequence for in vitro transcription. |

Supplementary table 9

**Supplementary table for Exo-C and reference NGS data SNV precision comparison.**

| Set of Golden Standard SNVs | Remaining reference dataset | SNV intersection percent with the reference dataset | SNV intersection percent with Exo-C dataset |
|---|---|---|---|
| (Dixon et al), (Moqin et al), (Wang et al) | (Ray et al) | 73,00% | 96,30% |
| (Dixon et al), (Moqin et al), (Ray et al) | (Wang et al) | 80,90% | 95,80% |
| (Dixon et al), (Wang et al), (Ray et al) | (Moqin et al) | 91,00% | 96,80% |
| (Wang et al), (Moqin et al), (Ray et al) | (Dixon et al) | 99,70% | 97,10% |

Golden standard SNVs were identified by overlapping SNV calls from three short-read datasets (column 1) and filtering for Exo-C capture
